# Functional delineation of tissue-resident CD8^+^ T cell heterogeneity during infection and cancer

**DOI:** 10.1101/2020.03.05.979146

**Authors:** J. Justin Milner, Clara Toma, Zhaoren He, Nadia S. Kurd, Quynh P. Nguyen, Bryan McDonald, Lauren Quezada, Christella E. Widjaja, Deborah A. Witherden, John T. Crowl, Gene W. Yeo, John T. Chang, Kyla D. Omilusik, Ananda W. Goldrath

## Abstract

Unremitting defense against diverse pathogens and malignancies requires a dynamic and durable immune response. Tissue-resident memory CD8^+^ T cells (Trm) afford robust protection against infection and cancer progression through continuous surveillance of non-lymphoid tissues. Here, we provide insight into how Trm confer potent and persistent immunity through partitioning of distinct cellular subsets differing in longevity, effector function, and multipotency. Antigen-specific CD8^+^ T cells localized to the epithelium of the small intestine are primarily comprised of a shorter-lived effector population most prominent early following both acute viral and bacterial infections, and a longer-lived Id3^hi^ Trm population that subsequently accumulates at later memory timepoints. We define regulatory gene-programs driving these distinct Trm states, and further clarify roles for Blimp1, T-bet, Id2, and Id3 in supporting and maintaining intestinal Trm heterogeneity during infection. Further, through single-cell RNAseq analysis we demonstrate that tumor-infiltrating lymphocytes broadly differentiate into discrete populations of short-lived and long-lived Trm-like subsets, which share qualities with terminally-exhausted and progenitor-exhausted cells, respectively. As the clinical relevance of Trm continues to widen from acute infections to settings of chronic inflammation and malignancy, clarification of the spectrum of phenotypic and functional states exhibited by CD8^+^ T cells that reside in non-lymphoid tissues will provide a framework for understanding their regulation and identity in diverse pathophysiological contexts.

## Introduction

Tissue-resident memory CD8^+^ T cells have emerged as critical mediators of health and disease. Trm remain permanently lodged in non-lymphoid tissues (and in some cases lymphoid tissues) without recirculating and are transcriptionally, epigenetically, functionally, and anatomically distinct from recirculating populations of CD8^+^ T cells (Masopust and Soerens, 2019; Milner and Goldrath, 2018; Szabo et al., 2019). The widespread relevance of Trm in preventing and precipitating disease stems from their unique localization patterns, robust inflammatory sentinel activity, potent effector function, and longevity at sites of imminent or ongoing immune responses (Ho and Kupper, 2019; Masopust and Soerens, 2019). Therefore, understanding the molecular signals controlling the fate, function, and homeostasis of Trm is relevant in diverse pathophysiological settings ranging from infection to cancer.

Upon infiltration into non-lymphoid sites, CD8^+^ T cells rapidly fine-tune epigenetic and gene-expression programs to adapt to diverse microenvironments, ultimately changing cellular metabolism, function, and tissue-retention (Milner and Goldrath, 2018). Dynamic environmental signals converge on several key Trm-fate-specifying transcription factors that modulate gene-expression programs controlling tissue-retention and egress. Blimp1, Hobit, and Runx3 repress the transcriptional signature associated with circulating memory cells and facilitate induction of a tissue-residency gene-expression program (Mackay et al., 2016; Milner and Goldrath, 2018). Conversely, T-bet, Eomes, and KLF2 can impede Trm formation (Laidlaw et al., 2014; Mackay et al., 2015; Skon et al., 2013). While it has become apparent that Trm are controlled by unique transcriptional signals, resolving Trm heterogeneity and the differentiation pathway of this unique cell type will contribute to a more integrated appreciation of transcription factor-mediated control of Trm fate and function.

The circulating CD8^+^ T cell population is heterogeneous, not only evolving over time but also composed of numerous subsets within infection timepoints (Arsenio et al., 2014; Kakaradov et al., 2017). The effector phase of infection predominantly consists of terminally-differentiated, short-lived KLRG1^hi^CD127^lo^ terminal effector (TE) cells and relatively fewer multipotent KLRG1^lo^CD127^hi^ memory-precursor (MP) cells (Chen et al., 2018). Some effector CD8^+^ T cells persist following infection at memory timepoints as long-lived effector cells (LLEC) (Olson et al., 2013; Omilusik et al., 2018), while others– predominately MP cells–continue to differentiate over time into long-lived, protective memory cells that can be broadly divided into central memory (Tcm) and effector memory (TEM) subsets. Tcm display enhanced lymphoid homing, multipotency, expansion potential, and longevity compared to Tem. Conversely, Tem are more terminally fated, shorter-lived, have limited expansion potential, and can elicit rapid effector function upon reinfection (Chen et al., 2018). This diversity in circulating memory T cell states endows the immune response with flexibility, allowing both rapid and sustained responses as well as the generation of secondary memory populations that retain these characteristics. Differentiation and maintenance of circulating memory subsets are also dynamically controlled by a compendium of transcription factors, including: Id2 (Cannarile et al., 2006; Knell et al., 2013; Masson et al., 2013), T-bet (Joshi et al., 2007), Blimp1 (Kallies et al., 2009; Rutishauser et al., 2009) Zeb2 (Dominguez et al., 2015; Omilusik et al., 2015), and STAT4 (Mollo et al., 2014) that are critical for Tem, and Id3 (Ji et al., 2011; Yang et al., 2011), Eomes (Banerjee et al., 2010; Pearce et al., 2003), Bcl6 (Ichii et al., 2002), Foxo1 (Hess Michelini et al., 2013; Rao et al., 2012), Tcf1 (Jeannet et al., 2010; Zhou et al., 2010), Zeb1 (Guan et al., 2018), Bach2 (Roychoudhuri et al., 2016), and STAT3 (Cui et al., 2011) that support Tcm differentiation. Trm heterogeneity has been alluded to in multiple non-lymphoid sites and infection models (Bergsbaken and Bevan, 2015; Bergsbaken et al., 2017; Boddupalli et al., 2016; Harrison et al., 2019; Kumar et al., 2018; Masopust and Soerens, 2019), and differing levels of CD69 and CD103 have been useful in studying Trm maturation. However it is unlikely these molecules capture the full spectrum of Trm heterogeneity, as Trm are often uniformly CD69^+^CD103^+^ in certain non-lymphoid sites such as the skin and small intestine or predominantly CD103^lo^ as in the kidney, heart and brain (Casey et al., 2012; Ma et al., 2017; Mackay et al., 2013). Thus, it remains unclear if the Trm population is comprised of distinct cell subsets with differing functional attributes and memory potential, analogous to circulating CD8^+^ T cells.

Facilitated by single-cell RNA-sequencing (scRNA-seq) analysis, we clarify Trm ontogeny and heterogeneity. Together with the companion paper (Kurd *et al*., submitted), we find significant inter- and intra-temporal molecular heterogeneity within the small intestine intraepithelial lymphocyte (siIEL) compartment that reveals distinct subsets not previously appreciated. Despite being transcriptionally distinct, siIEL CD8^+^ T cell subsets share qualities with the circulating CD8^+^ T cell response, and can be phenotypically, transcriptionally, and functionally distinguished based on the reciprocal expression of the effector and memory-associated transcriptional regulators, Blimp1 and Id3. These studies provide a framework to better understand the transcriptional signals controlling Trm differentiation and longevity, especially as Trm continue to be explored in diverse disease states beyond acute infections. In connection, we extend our findings to the tumor context, highlighting evidence of distinct Trm-like subsets within the tumor microenvironment.

## Results

### Anti-viral siIEL CD8^+^ T cells are inter- and intra-temporally heterogeneous

Trm in all non-lymphoid tissues examined to date are transcriptionally distinct from circulating memory populations (Mackay et al., 2013; Milner et al., 2017; Wakim et al., 2012). To expand on these findings, we profiled the transcriptome of Tcm (CD62L^+^) and Tem (CD62L^−^) from the spleen as well as siIEL Trm isolated >50 days following acute LCMV infection. Consistent with previous reports, Trm were transcriptionally distinct from Tcm and Tem based on hierarchical clustering analysis, (Figure 1A) and enriched with a core tissue-residency gene-expression signature (Milner et al., 2017) (Figure 1B). However, we identified four gene expression modules (clusters 2-5) commonly regulated between Trm and Tcm or Trm and Tem, indicating that Trm integrate Tcm and Tem characteristics as well. Furthermore, Trm were found to exhibit mixed expression of both Tcm and Tem fate-specifying transcription factors (Figure 1A). These data highlight that although Trm comprise a distinct memory T cell subset, Trm also share certain transcriptional features with the more effector-like Tem cells as well as the longer-lived Tcm population, consistent with findings from Kurd *et al*. (submitted). To further clarify the Trm population in terms of a long-lived memory or effector phenotype, we assessed expression of gene signatures in a triwise comparison of Tcm, Tem, and Trm populations (van de Laar et al., 2016). We found, as expected, that Tcm displayed enrichment of a long-lived memory CD8^+^ T cell gene-signature relative to Tem (Figure 1B). However, some of these memory-associated genes were also upregulated in Trm relative to both Tem and Tcm. Conversely, Trm and Tem exhibited greater expression of effector-associated genes compared to Tcm, but Trm also displayed elevated expression of numerous effector-associated genes compared to Tem. Therefore, the non-mutually exclusive possibilities arise—does the Trm population exist in a unique state simultaneously balancing both effector and memory qualities, or is it comprised of distinct effector- and memory-like subsets undetectable via bulk sequencing analyses?

**Figure 1.**
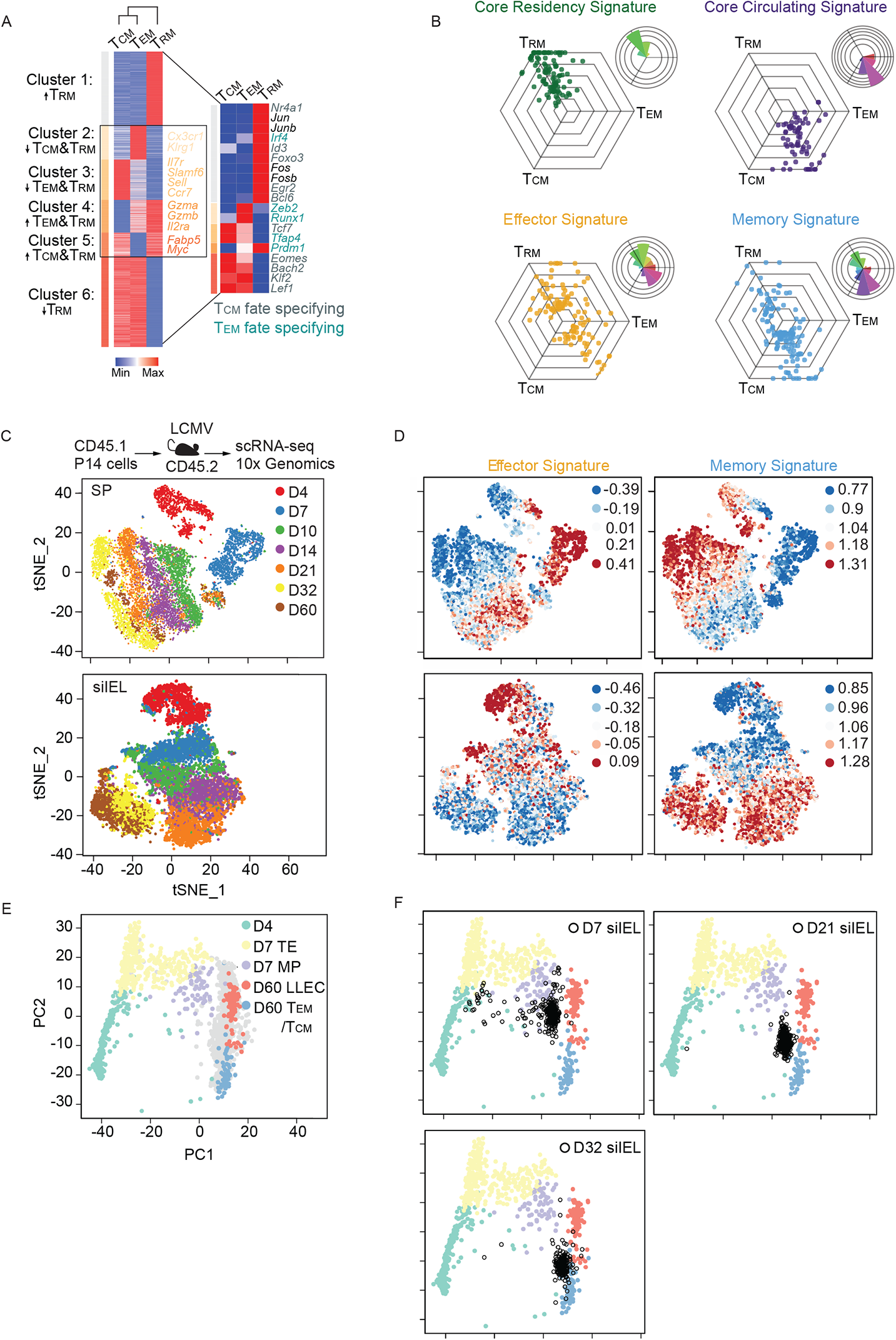
The anti-viral tissue-resident siIEL CD8^+^ T cell population is heterogeneous. P14 CD8^+^ T cells were transferred into congenically distinct hosts that were subsequently infected with LCMV. Donor cells from the spleen and siIEL were sorted over the course of infection for bullk RNA-seq or scRNA-seq. **(A)** Heatmap illustrating the relative expression of genes differentially expressed among Tcm, Tem and Trm populations from bulk RNA-seq analysis; gene clusters are ordered through K-means clustering analysis. Transcriptional regulators reported as important for Tcm (gray) and Tem (teal) fate are highlighted (right). **(B**) To visualize relative expression levels in a triwise comparison (van de Laar et al., 2016), filtered genes differentially expressed and present within the designated gene list were plotted in a hexagonal diagram in which distance of an individual data point represents gene expression enrichment or depletion, and the magnitude of upregulation is reflected by the distance from the origin. Rose plots (upper right corner of each hexagonal plot) indicate the percentages of genes in each orientation. Genes of the core residency (green), core circulating (purple), effector (yellow), and memory (blue) signatures are highlighted. **(C-F)** scRNA-seq analysis across an infection time course. tSNE plots of cells from the siIEL or spleen (SP) over all infection timepoints **(C)** colored by indicated timepoint or **(D)** colored by intensity of effector or memory gene signatures. **(E)** Principal component analysis of the spleen samples according to the expression of the differentially expressed genes between effector and memory cells. Day 4 (green), day 7 TE (yellow) or MP (purple), and day 60 LLEC (pink) or Tem/Tcm (blue) are highlighted while samples from remaining timepoints are shaded grey. **(F)** The siIEL CD8^+^ T cell samples (black) are projected onto the 2D space according to the same principal components.

To investigate the apparent dual effector-memory state of the Trm population, we utilized scRNA-seq profiling of LCMV GP_33-41_-specific P14 CD8^+^ T cells isolated from the spleen or siIEL compartment over the course of an LCMV infection (Figure 1C). Data from all timepoints from the spleen or siIEL were integrated into an unsupervised t-distributed stochastic neighborhood embedding (tSNE) analysis (Figure 1C-D). Similar to circulating CD8^+^ T cells from the spleen, siIEL CD8^+^ T cells were enriched for an effector gene signature early after infection, and subsequently, were enriched for a memory gene signature at later infection timepoints (Figure 1D). Thus, the siIEL CD8^+^ T cells exhibited substantial inter-temporal heterogeneity. We also detected intra-temporal heterogeneity within the siIEL CD8^+^ T cell population at both early and late infection timepoints. siIEL CD8^+^ T cells enriched with a memory gene signature could be observed as early as day 4 following infection, and those with an enrichment of the effector gene signature were evident on days 32 and 60 of infection. Furthermore, few CD8^+^ T cells appear to simultaneously display enrichment of both effector and memory gene signatures (Figure 1D). To orient the transcriptional profile of siIEL CD8^+^ T cells to that of the splenic populations, principal component analysis was performed on all splenic scRNA-seq samples according to the expression of the differentially expressed genes between effector and memory T cell states. All splenic cells were plotted according to the top two principal components and day 4 effector T cells, day 7 TE and MP, and day 60 LLEC and Tem/Tcm were highlighted (Figure 1E). Here, the bulk CD62L^−^ population was subdivided into CD127^−^ and CD127^+^ cells, which resolves LLEC and Tem subsets, allowing a more refined delineation of effector and memory gene-expression signatures. Next, siIEL CD8^+^ T cells from the indicated time points of infection were projected into the 2D space according to the same PCs (Figure 1F; black). At day 7 of infection, siIEL CD8^+^ T cells clustered near the circulating effector T cell subsets and the LLEC. However, by day 21 of infection, the siIEL CD8^+^ T cells are interposed between LLEC and Tem/Tcm populations. Therefore, Figure 1F demonstrates that Trm are transcriptionally distinct from circulating memory populations, but Trm exist in a range of effector and memory states analogous to circulating memory CD8^+^ T cells. In summary, we find that siIEL CD8^+^ T cells exhibit both inter- and intra-temporal heterogeneity, and the current broad definition of Trm is not necessarily reflective of this apparent diversity as effector-like siIEL CD8^+^ T cells also persist into the memory phase of infection.

### Key regulatory molecules distinguish effector and memory subsets of siIEL CD8^+^ T cells

As we have noted distinct sub-populations of Trm, somewhat analogous to circulating CD8^+^ T cell subsets, we next sought to evaluate how the current paradigm of circulating CD8^+^ T cell diversity parallels Trm heterogeneity. We first assessed the inter-temporal expression dynamics of molecules often utilized to clarify heterogeneity within the circulating CD8^+^ T cell compartment; as anticipated, circulating CD8^+^ T cells exhibited an inverse expression pattern of *Klrg1* and *Il7r* (encoding CD127, Figure 2A). Despite inter-temporal heterogeneity in effector and memory gene signatures within the siIEL CD8^+^ T cell populations (Figure 1D), *Il7r* and *Klrg1* expression kinetics were not as dynamic as in splenic CD8^+^ T cells (Figure 2A). Nonetheless, *Klrg1* expression was elevated early (days 4-10) in the intestine and diminished over time, whereas *Il7r* expression steadily increased. This gene-expression pattern is similarly reflected at the protein level (Figure 2B). The expression of pro-effector transcriptional regulators *Tbx21* (encoding T-bet)*, Prdm1* (encoding Blimp1)*, Zeb*2, and *Id2* as well as pro-memory factors *Bcl6, Eomes, Tcf7* (encoding TCF1), and *Id3* were assessed across all siIEL samples (Figure 2C). Consistent with Figure 1A, the expression pattern of pro-memory or pro-effector transcriptional regulators did not conform to an expected expression pattern (i.e. pro-effector and pro-memory transcriptional regulators being upregulated at early and later infection timepoints, respectively). *Prdm1* and *Eomes* expression peaked early in siIEL CD8^+^ T cells whereas *Id3* and *Bcl6* expression increased over time. Within siIEL CD8^+^ T cells, *Tbx21, Tcf7*, and *Id2* expression levels remained relatively uniform, although *Id2* expression was elevated at day 7 of infection and then subtly decreased (Figure 2C). Intracellular staining for T-bet, Eomes, and TCF1 in siIEL CD8^+^ T cells following LCMV infection was reflective of the scRNA-seq results (Figure 2E). Therefore, differential KLRG1 and CD127 expression indicate siIEL CD8^+^ T cell heterogeneity, and the expression pattern of select transcription factors may be indicative of effector and memory siIEL CD8^+^ cell states.

**Figure 2.**
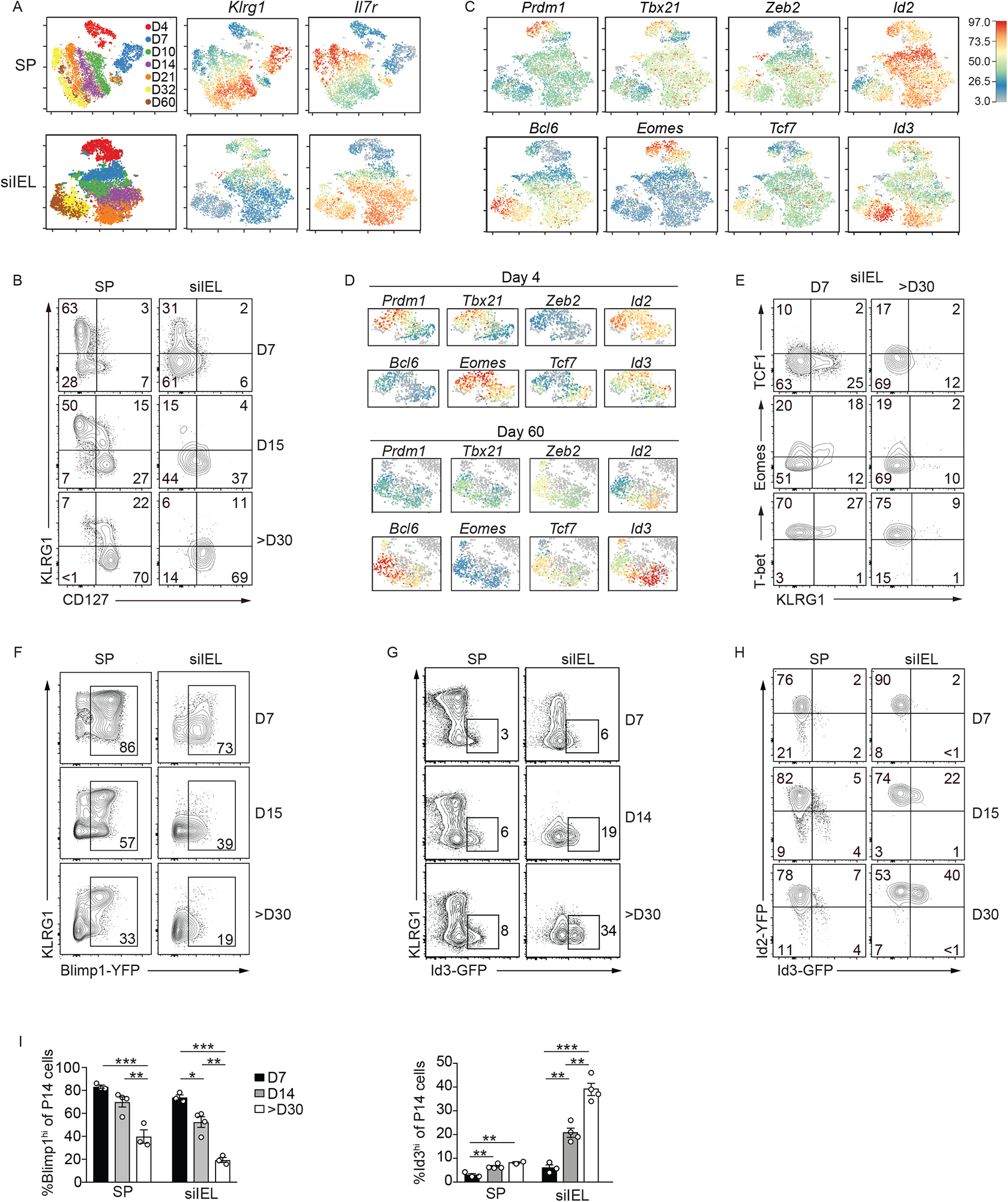
Heterogeneous expression of key regulatory factors by siIEL CD8^+^ T cells. P14 CD8^+^ T cells were transferred into congenically distinct hosts that were subsequently infected with LCMV. Donor cells from the spleen and siIEL were sorted over the course of infection for scRNA-seq or were analyzed through flow cytometry. **(A)** tSNE plots of cells from the spleen (top) or siIEL (bottom) over all infection timepoints colored by sample (left) and intensity of *Klrg1* (middle) or I*l7r* (right) mRNA levels. **(B)** Expression levels of CD127 and KLRG1 during LCMV infection. **(C)** Relative gene-expression of highlighted transcriptional regulators in siIEL cells following the map from A. **(D)** Expression of indicated transcriptional regulators within siIEL CD8^+^ T cells at day 4 (top) or 60 (bottom) of infection. **(E)** Expression levels of indicated transcriptional regulators. **(F-H)** Blimp1-YFP, Id3-GFP or Id2-YFP/Id3-GFP P14 CD8^+^ T cells were transferred into congenically distinct hosts that were infected with LCMV. At indicated times of infection, reporter expression was analyzed in the spleen and siIEL by flow cytometry. **(I)** The percentage of Blimp1-YFP^hi^ and Id3-GFP^hi^ P14 cells in the spleen and siIEL are quantified. Numbers in plots represent the frequency of cells in the indicated gate. All data are from one representative experiment of 2 independent experiments with n=3-5 (B,E,H) or n=2-4 (F,G). Graphs show mean ± SEM; *p<0.05, **p<0.01, ***p<0.001.

We noted intra-temporal siIEL CD8^+^ T cell heterogeneity at both effector and memory phases of infection (Figure 2D), similar to the companion study by Kurd *et al*. (submitted). For example, siIEL CD8^+^ T cells with high and low expression of *Prdm1*, *Tbx21*, and *Eomes* or *Zeb2* and *Bcl6* were noted among cells at early (day 4) or later (day 60) timepoints of infection, respectively. Several transcriptional regulators (*Id2*, *Tcf7* and *Id3*) were differentially expressed among individual cells within the siIEL CD8^+^ T cell population throughout LCMV infection. Two notable transcriptional regulators with dynamic expression over the course of infection were *Prdm1* and *Id3*. *Prdm1* expression was highest in siIEL CD8^+^ T cells at day 4 of infection then declined over time (Figure 2C), consistent with the *Prdm1* expression patterns reported for skin Trm (Mackay et al., 2016). To extend this to siIEL Trm and the observed population heterogeneity, we utilized Blimp1-YFP-expressing P14 CD8^+^ T cells to assess Blimp1 expression in siIEL CD8^+^ T cells over the course of LCMV infection. Consistent with the scRNA-seq profiling, Blimp1 expression was highest early following infection and corresponded with KLRG1 expression but decreased over time (Figure 2F). Within the circulation, Id3 expression marks CD8^+^ T cells with greater memory potential (Ji et al., 2011; Yang et al., 2011), but its function in tissue-resident populations has yet to be clarified. We next used Id3-GFP-reporter (Miyazaki et al., 2011) P14 CD8^+^ T cells to evaluate Id3 expression in siIEL CD8^+^ T cells over the course of LCMV infection. Similar to that observed in circulating CD8^+^ T cells (Ji et al., 2011) and consistent with our scRNA-seq data, Id3 expression inversely correlated with that of Blimp1 and the frequency of KLRG1^lo^Id3^hi^ cells increased over time wherein nearly one-third of all Trm within the siIEL compartment at day ~30 of infection expressed Id3 (Figure 2G). Notably, the dramatically increased frequency of Id3-expressing was cells not observed in the circulating memory population, which peaked at ~8%. Id2 has a recognized role in promoting survival and terminal differentiation of circulating CD8^+^ T cells (Cannarile et al., 2006; Knell et al., 2013; Masson et al., 2013; Omilusik et al., 2018). Therefore, we utilized Id2-YFP-expressing P14 CD8^+^ T cells (Yang et al., 2011), and found that Id2 expression was slightly upregulated compared to splenic CD8^+^ T cells, but as observed in splenic populations (Yang et al., 2011), expression levels remained relatively constant over time (Figure 2H and Figure S2B). Taken together, siIEL CD8^+^ T cells at effector timepoints expressed elevated levels of Blimp1 and KLRG1, whereas Id3 was reciprocally expressed in siIEL CD8^+^ T cells and continually increased into the memory phase of infection.

### Blimp1 and Id3 delineate distinct populations of siIEL CD8^+^ T cells

To examine the relationship between Blimp1 and Id3 expression in the context of the observed siIEL CD8^+^ T cell heterogeneity, we generated P14 Blimp1-YFP/Id3-GFP-reporter mice. In all tissues examined (Figure 3A,C), Blimp1 and Id3 expression was mutually exclusive, consistent with the reported function of Blimp1 as a transcriptional repressor of Id3 expression (Ji et al., 2011). Following LCMV infection, the proportion of Blimp1^hi^ CD8^+^ T cells decreased over time across all tissues examined. In the spleen and mesenteric lymph node, Id3 expression remained relatively unchanged past day 14 of infection; however, the frequency of Id3-expressing siIEL CD8^+^ T cells increased over time, with a ~10-fold increase in the percentage of Id3^hi^ siIEL CD8^+^ T cells from day 7 to day 80, and a corresponding ~50-fold loss in the absolute number of Blimp1-YFP siIEL CD8^+^ T cells (Figure 3A,B). We next evaluated if tissue-resident cells in other non-lymphoid microenvironments exhibited similar expression patterns of Blimp1 and Id3 (Figure 3C). Interestingly, we found that CD8^+^ T cell populations were transcriptionally varied across non-lymphoid sites, wherein only siIEL and lamina propria compartments had a sizeable Id3^hi^ population of CD8^+^ T cells at day 90 of LCMV infection. Kidney and salivary gland contained a low frequency of Id3^hi^ CD8^+^ T cells while white adipose tissue and brain populations had no detectable Id3. We next evaluated Id3 expression patterns in epidermal Trm in a model of skin inflammation and detected robust expression of Id3 in transferred CD8^+^ T cells by day 14 following DNFB treatment (Figure 3D). Thus, while Blimp1 is expressed in a subset of CD8^+^ T cells across multiple non-lymphoid sites, Id3 expression may be restricted to CD8^+^ T cells within barrier tissues. We also found similar Blimp1 and Id3 expression patterns in siIEL CD8^+^ T cells following enteric *Listeria monocytogenes* infection (Figure 3E). Therefore, Id3 and Blimp1 expression were dynamically and inversely regulated within CD8^+^ T cell populations resident to barrier tissues in the context of infection and inflammation.

**Figure 3.**
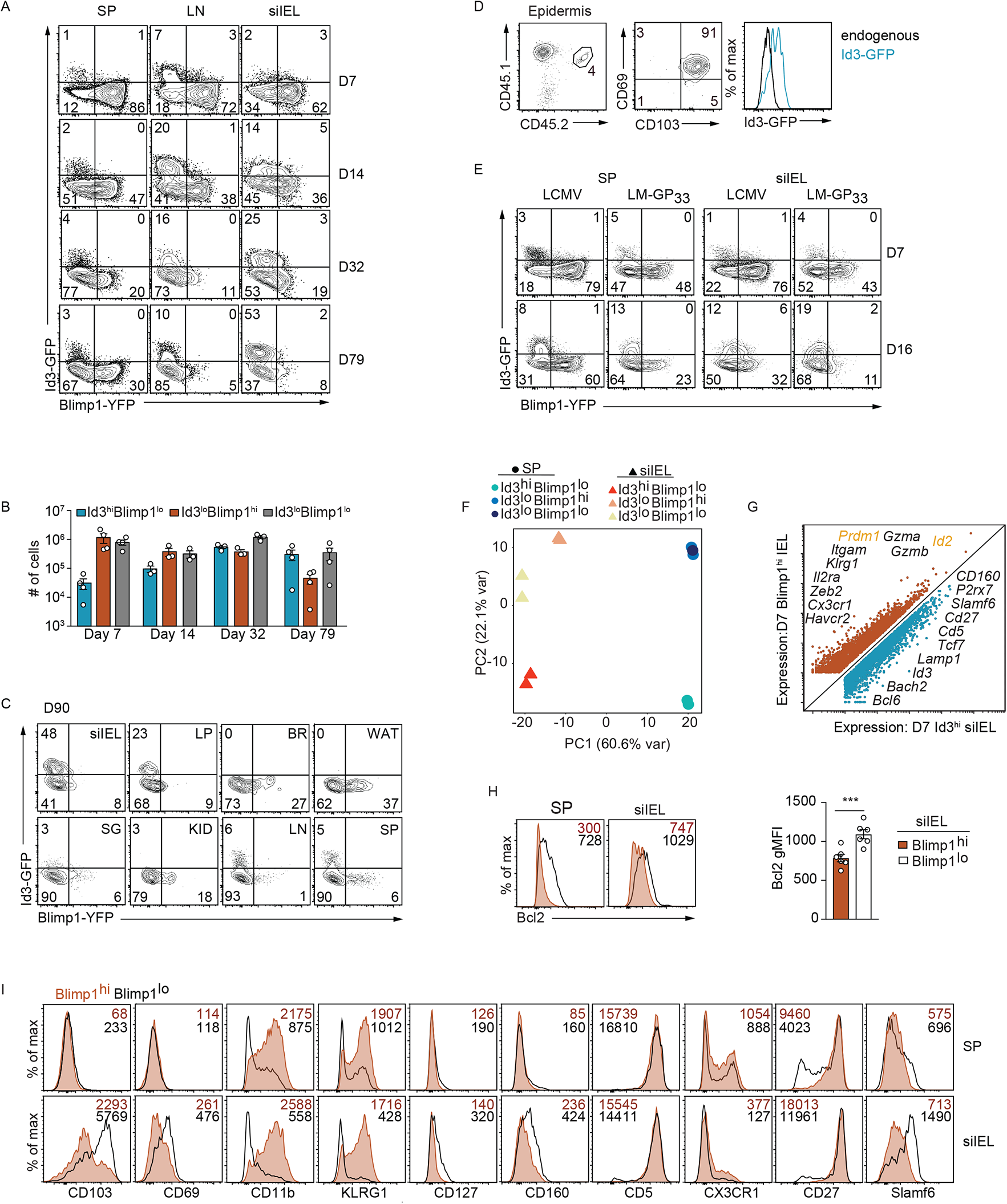
Blimp1 and Id3 reporter expression identifies distinct subsets of siIEL CD8^+^ T cells. Congenically distinct Id3-GFP or Blimp1-YFP/Id3-GFP (double reporter) P14 CD8^+^ T cells were transferred to wild-type hosts that were subsequently infected with LCMV i.p. or LM-GP_33_ o.g. **(A)** Blimp1-YFP and Id3-GFP reporter expression was assessed by flow cytometry in double reporter P14 cells from indicated host tissue over the course of the LCMV infection. **(B)** Quantification of the number of cells in the indicated populations from A. **(C)** Blimp1-YFP and Id3-GFP reporter expression in double reporter P14 cells from indicated host tissues on day 90 of LCMV infection is shown. **(D)** *In vitro* activated Id3-GFP P14 CD8^+^ T cells were adoptively transferred into recipient mice treated with DNFB on the left flank. Within the epidermis, the frequency of transferred cells among CD8^+^ T cells (left) and corresponding CD69, CD103 (middle), and Id3-GFP expression (right) on day 14 following treatment is indicated. **(E)** Blimp1-YFP and Id3-GFP reporter expression in double reporter P14 CD8^+^ T cells from host spleen and siIEL on days 7 and 16 following LCMV and LM-GP_33_ infection is compared. **(F-G)** On day 7 of infection, Id3^hi^Blimp1^lo^, Id3^lo^Blimp1^hi^, and Id3^lo^Blimp1^lo^ P14 CD8^+^ T cells from the spleen and siIEL were sorted for RNA-sequencing. Principal component analysis (F) of gene expression from the sorted P14 CD8^+^ T cell populations is shown, or differentially expressed genes (G) between Id3^hi^Blimp1^lo^ and Id3^lo^Blimp1^hi^ siIEL P14 CD8^+^ T cells are highlighted. **(H-I)** Blimp-YFP or Blimp1-YFP/Id3-GFP P14 CD8^+^ T cells transferred into congenically distinct hosts that were infected with LCMV were isolated from spleen and siIEL and analyzed on day 7 of infection for expression of Bcl2 (H) or indicated surface molecules (I). siIEL = small intestine intraepithelial lymphocytes; LP = small intestine lamina propria; BR= brain; WAT= white adipose tissue; SG = salivary gland; KID = kidney; LN = mesenteric lymph node; SP= spleen. Numbers in plots represent the frequency (A,C,D,E) or gMFI (H,I) of cells in the indicated gate. All data are from one representative experiment of 2-3 independent experiments with n=3-5 (A-G,I) or cumulative from 2 independent experiments with a total n=6 (H). Analysis of Slamf6 expression (D) is one independent experiment, n=3.

Although the frequency of Id3-expressing cells continuously increased over time, we observed a small proportion of siIEL CD8^+^ T cells that expressed Id3 at the peak of LCMV infection (Figure 3A), suggesting that subset heterogeneity may be established early after infection. Principal component analysis of gene expression by Id3^lo^Blimp^hi^, Id3^hi^Blimp1^lo^, and Id3^lo^Blimp1^lo^ siIEL and splenic CD8^+^ T cells on day 7 of LCMV infection revealed that effector CD8^+^ T cells cluster based on tissue origin, but also that variation between the three siIEL populations is evident even at early effector timepoints (Figure 3F). Comparison of gene expression between the Id3^lo^Blimp^hi^ and Id3^hi^Blimp1^lo^ siIEL CD8^+^ T cell subsets highlighted that considerable transcriptional differences already existed by day 7 of infection, with Blimp1^hi^ cells expressing canonical effector molecules such as *Cx3cr1*, *Zeb2*, *Klrg1*, *Id2*, and *Gzma/b* whereas Id3^hi^ cells expressed canonical memory genes including *Bcl6*, *Bach2*, *Tcf7, and Cd27* (Figure 3G), suggesting this population includes the long-lived Trm precursor cells. This early subset variation was further confirmed by correlating levels of key molecules (KLRG1, CX3CR1, CD27, Slamf6, and CD69) with Blimp1-YFP expression by siIEL CD8^+^ T cells at day 7 of infection (Figure 3H-I). Thus, as early as day 7 of infection, siIEL CD8^+^ T cells were transcriptionally distinct from circulating effector populations and exhibited considerable subset heterogeneity, reminiscent of circulating TE and MP populations.

### Id3^hi^ and Blimp1^hi^ siIEL CD8^+^ T cells distinguish functional Trm subsets

To determine if the expression pattern of Blimp1 and Id3 observed within the siIEL CD8^+^ T cell population reflects distinct subsets of non-lymphoid tissue T cells, we next profiled the transcriptome of the Id3- and Blimp1-expressing siIEL CD8^+^ T cell subpopulations on day 35 of infection. As expected, principal component analysis revealed that, despite differential expression of Blimp1 and Id3, all three populations sorted from the siIEL compartment were transcriptionally similar relative to the corresponding circulating memory CD8^+^ T cells within the spleen and lymph node (Figure 4A). However, subtle differences between the siIEL populations were detectable in this context (Figure 4A), and closer examination revealed >2000 differentially expressed transcripts between Blimp1^hi^Id3^lo^ and Blimp1^lo^Id3^hi^ siIEL CD8^+^ T cells (Figure 4B), including effector-associated genes (*Klrg1*, *Zeb2*, *Bhlhe40* and *Gzma/b*) upregulated in the Blimp1^hi^Id3^lo^ subset and memory-associated genes (*Tcf7*, *Eomes*, *Bach2*) upregulated in the Blimp1^lo^Id3^hi^ population. Flow cytometry analysis of siIEL CD8^+^ T cells on days 22-25 of infection confirmed many of the findings from the RNA-seq analysis (Figure 4C). Thus, while the siIEL Trm population as a whole was transcriptionally distinct from the circulating memory CD8^+^ T cell populations, Blimp1 and Id3 delineated distinct subsets.

**Figure 4.**
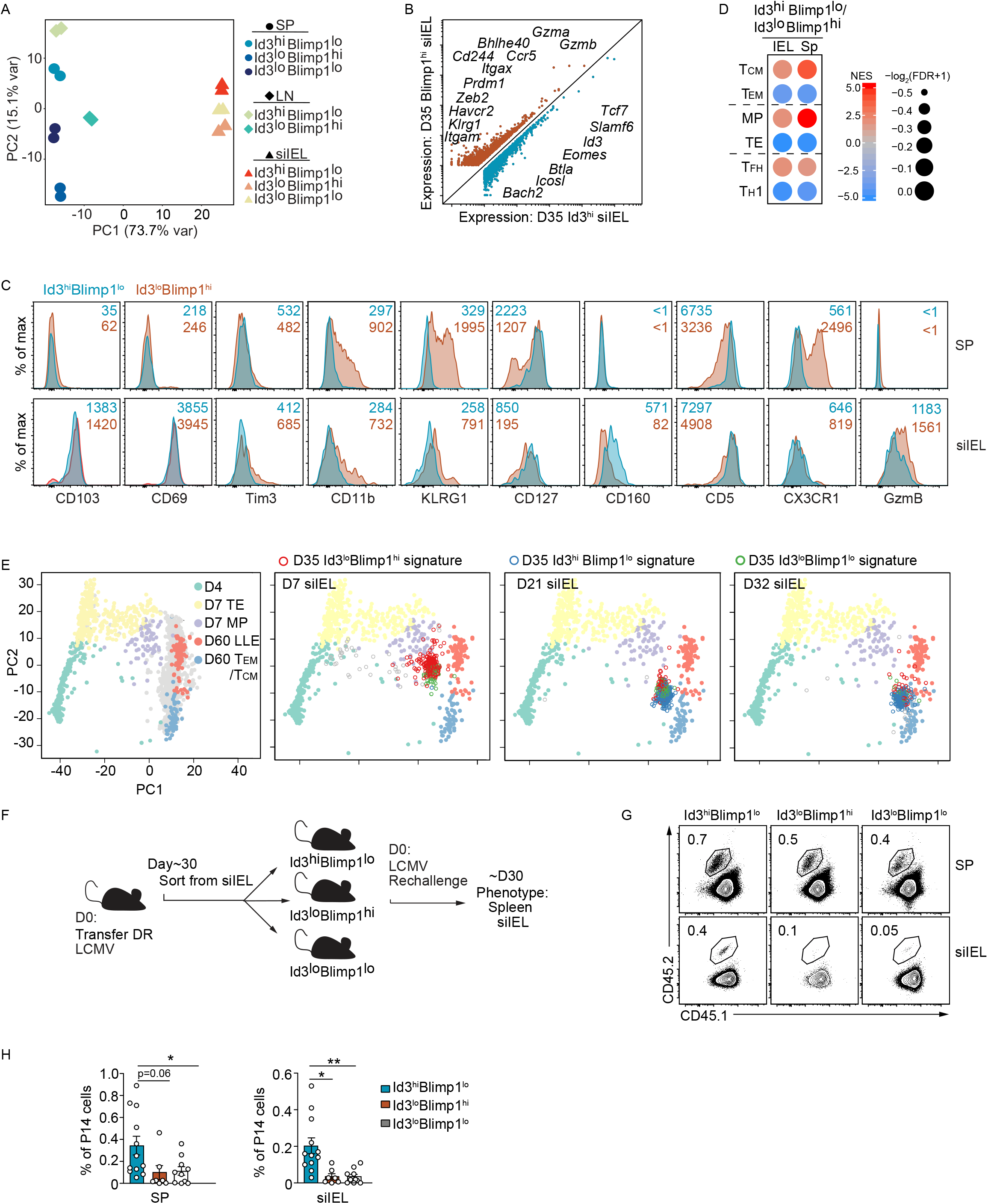
Id3^hi^ siIEL CD8^+^ T cells exhibit a long-lived memory state relative to the Blimp1^hi^ subset. **(A-D)** Blimp1-YFP/Id3-GFP P14 CD8^+^ T cells were transferred into congenically distinct hosts that were subsequently infected with LCMV. On day 35 of infection, Id3^hi^Blimp1^lo^, Id3^lo^Blimp1^hi^, and Id3^lo^Blimp1^lo^ P14 CD8^+^ T cells from the spleen (SP), mesenteric lymph node (LN), and siIEL were sorted for RNA-sequencing. **(A)** Principal component analysis of gene expression from the sorted P14 CD8^+^ T cell populations is shown. **(B)** Genes differentially expressed by Id3^hi^Blimp1^lo^ and Id3^lo^Blimp1^hi^ P14 siIEL CD8^+^ T cells are highlighted. **(C)** Blimp1-YFP/Id3-GFP P14 CD8^+^ T cells transferred into congenically distinct hosts subsequently infected with LCMV were analyzed in spleen and siIEL on day 25 of infection. Expression of indicated molecules were compared between Id3^hi^Blimp1^lo^ (teal) and Id3^lo^Blimp1^hi^ (orange) subsets. Numbers in plots represent gMFI. **(D)** GSEA for specified gene signatures enriched in Id3^hi^Blimp1^lo^ and Id3^lo^Blimp1^hi^ siIEL CD8^+^ T cell subsets. **(E)** Principal component analysis (from scRNA-seq dataset) of the spleen samples according to the expression of the differentially expressed genes between effector and memory cells. Day 4 (green), day 7 TE (yellow) or MP (purple), and day 60 LLE (pink) or Tem/Tcm (blue) are highlighted. The siIEL CD8^+^ T cells samples are projected to the 2D space according to the same principal components and enrichment for Id3^hi^Blimp1^lo^ (blue), Id3^lo^Blimp1^hi^ (red), and Id3^lo^Blimp1^lo^ (green) gene signatures is indicated. **(F)** Schematic of experimental set-up. Congenically distinct Blimp1-YFP/Id3-GFP P14 CD8^+^ T cells were transferred to wild-type hosts that were infected with LCMV. More than 30 days after infection, Id3^hi^Blimp1^lo^, Id3^lo^Blimp1^hi^, and Id3^lo^Blimp1^lo^ P14 CD8^+^ T cells were sorted from the siIEL, and then retransferred intravenously into congenically distinct hosts subsequently infected with LCMV. After 30 days of infection, donor cells in the host spleen (SP) and siIEL were analyzed by flow cytometry. **(G)** Frequency of transferred cells among CD8^+^ T cells is shown. Numbers in plots represent the frequency of cells in the indicated gate. **(H)** Quantification of populations from G. All data are from one representative experiment of 2 independent experiments with n=3-4 (C) or representative (G) or cumulative (H) of 4 independent experiments with a total n=7-12. Graphs show mean ± SEM; *p<0.05, **p<0.01.

To place these Trm subsets in the context of other conventional T cell populations, we performed gene set enrichment analysis (GSEA). The gene-expression signature of Blimp1^lo^Id3^hi^ siIEL CD8^+^ T cells was enriched in genes associated with Tcm, MP and Tfh (i.e. cells that exhibit increased proliferation potential or memory-like qualities). In contrast, the Blimp1^hi^Id3^lo^ gene-expression signatures was enriched in genes associated with Tem, TE, and CD4^+^ TH1 (i.e. cells that are more terminally differentiated and exhibit effector-like properties) (Chen et al., 2018; Nguyen et al., 2019) (Figure 4D). Further, the siIEL CD8^+^ T cell subsets were next examined within the context of the principal component analysis performed in Figure 1E; gene-expression signatures of the distinct Id3- and Blimp1-expressing subsets were projected onto the 2D space with the top 2 principal components (Figure 4E). As expected, siIEL CD8^+^ T cells on day 7 of infection were enriched for the Id3^lo^Blimp1^hi^ signature (red) while the population at day 32 of infection was enriched with the Id3^hi^Blimp1^lo^ gene-expression signature (blue). Notably, the day 21 siIEL CD8^+^ T cell population was heterogeneous, comprised of cells clustering between the LLEC and Tem/Tcm populations and exhibiting differential enrichment of the distinct Id3/Blimp1 gene-expression signatures, revealing individual cells enriched for the different signatures.

To determine if Blimp1 and Id3 expression defined siIEL CD8^+^ T cell subsets with distinct memory potential, Blimp1^hi^Id3^lo^, Blimp1^lo^Id3^hi^ and Blimp1^lo^Id3^lo^ siIEL CD8^+^ T cell populations were sorted and transferred into recipient mice that were challenged with LCMV. The donor cell populations were analyzed on day 30 of infection (Figure 4F). Notably, we found that Blimp1^lo^Id3^hi^ IEL donor cells yielded a greater frequency of both circulating and resident populations following rechallenge compared to the subsets that did not express Id3 (Figure 4G,H). Thus, Id3^hi^ siIEL CD8^+^ T cells possessed enhanced proliferative capacity and multipotency as they formed a larger pool of memory cells and were capable of giving rise to both circulating and Trm populations, indicating that Id3 expression marks Trm with secondary memory potential.

### Transcriptional regulation of siIEL CD8^+^ T cell heterogeneity

We next assessed the role of key transcriptional regulators, associated with promoting TE versus MP fates, in driving the heterogeneity of siIEL CD8^+^ T cells. *Prdm1*^fl/fl^-Gzmb-Cre^+^ or *Id2*^f/f^-CD4-Cre^+^ P14 CD8^+^ T cells were mixed 1:1 with congenically distinct WT P14 CD8^+^ T cells and transferred to naive recipient mice subsequently infected with LCMV (Figure S3A,B,D,E). We previously noted that KLRG1 followed a relatively similar expression trajectory as Blimp1 and that Blimp1^hi^ cells expressed elevated levels of KLRG1 (Figure 2F). Therefore, we utilized KLRG1 to signify Blimp^hi^ siIEL CD8^+^ T cells in order to understand the roles of Blimp1 and Id2 in regulating the fate of distinct siIEL CD8^+^ T cell subsets in the absence of the reporters. On day 7 or 8 of infection, Blimp1 was generally required for optimal formation of bulk siIEL CD8^+^ T cells as previously reported (Mackay et al., 2016), but notably, the KLRG1^hi^ effector subset was more dramatically impacted by loss of Blimp1 (Figure S3D,E). Further, Id2-deficiency resulted in a selective loss of the KLRG1^hi^ siIEL CD8^+^ T cells, but interestingly the formation of the KLRG1^lo^ subset was not impacted (Figure S3D,E). We also examined if T-bet regulated the KLRG1^hi^Blimp1^hi^ siIEL CD8^+^ T subset by comparing siIEL *Tbx21* ^+/+^ and *Tbx21*^+/−^ P14 T cells in a similar mixed transfer experiment (Figure S3C-E). We found that T-bet also supported the formation of KLRG1^hi^ siIEL CD8^+^ T cells, but was not essential for the KLRG1^lo^ Trm precursor population in the siIEL compartment. Further, *Id2*-deficiency or *Tbx21*-heterozygosity resulted in elevated expression levels of CD103, CD69, Slamf6, CD27, and CD127 as well as lower expression levels of KLRG1 and CX3CR1 (Figure S3F). Taken together, the effector-associated transcriptional regulators Blimp1, Id2 and T-bet direct formation of the shorter-lived KLRG1/Blimp^hi^ siEL CD8^+^ T cell subset.

Given the unique expression pattern of Id3 and its role in supporting long-lived circulating memory cells, we examined Id3-mediated regulation of Trm maintenance by inducing deletion in established Trm cells. *Id3*^f/f^ -ERCre^+^ P14 CD8^+^ T cells were mixed 1:1 with *Id3*^+/+^ P14 CD8^+^ T cells and transferred to congenically distinct naive mice that were subsequently infected with LCMV. More than 30 days after infection, deletion was induced by treatment with tamoxifen. More than thirty days post-tamoxifen treatment, we assessed how induced deletion of Id3 impacted the homeostasis and phenotype of the Trm population (Figure 5A-B). We hypothesized that Id3 would be specifically required for maintenance of the long-lived Id3^hi^CD127^hi^ population of Trm, with less of a role in maintaining the effector-like Blimp1^hi^CD127^lo^ population. However, induced deletion of *Id3* resulted in a minor loss of Trm, and both CD127^hi^ and CD127^lo^ populations were equivalently impacted. We speculated the minimal phenotype observed by induced deletion of *Id3* was due to compensation by Id2, as we find *Id2* is abundantly expressed in both CD127^hi^ and CD127^lo^ Trm (Figure 2H). Utilizing a similar induced deletion system, we tested the impact of *Id2* deletion alone in maintaining the distinct Trm populations (Figure 5C). Here, we found that induced deletion of *Id2* in established Trm dramatically impacted the CD127^hi^ long-lived Trm population while minimally impacting the maintenance of the CD127^lo^ subset. Finally, *Id2*^f/f^*Id3*^f/f^-ERCre^+^ P14 cells were studied to allow simultaneous deletion of both *Id2* and *Id3* in established Trm cells (Figure 5D). Notably, deficiency of both *Id2* and *Id3* resulted in an even greater loss of siIEL CD8^+^ T cells than deletion of *Id2* alone (~4-fold greater loss of Trm with double deficiency compared to *Id2* deficiency), wherein the CD127^hi^ siIEL CD8^+^ T cell population was more profoundly affected than the CD127^lo^ subset (Figure 5E). Further, dual deficiency of *Id2* and *Id3* enhanced the frequency of CD11b^+^Tim3^+^ cells (Figure 5F). Thus, Id2 and Id3 are critical regulators of Trm maintenance, especially for long-lived CD127^hi^ Trm.

**Figure 5.**
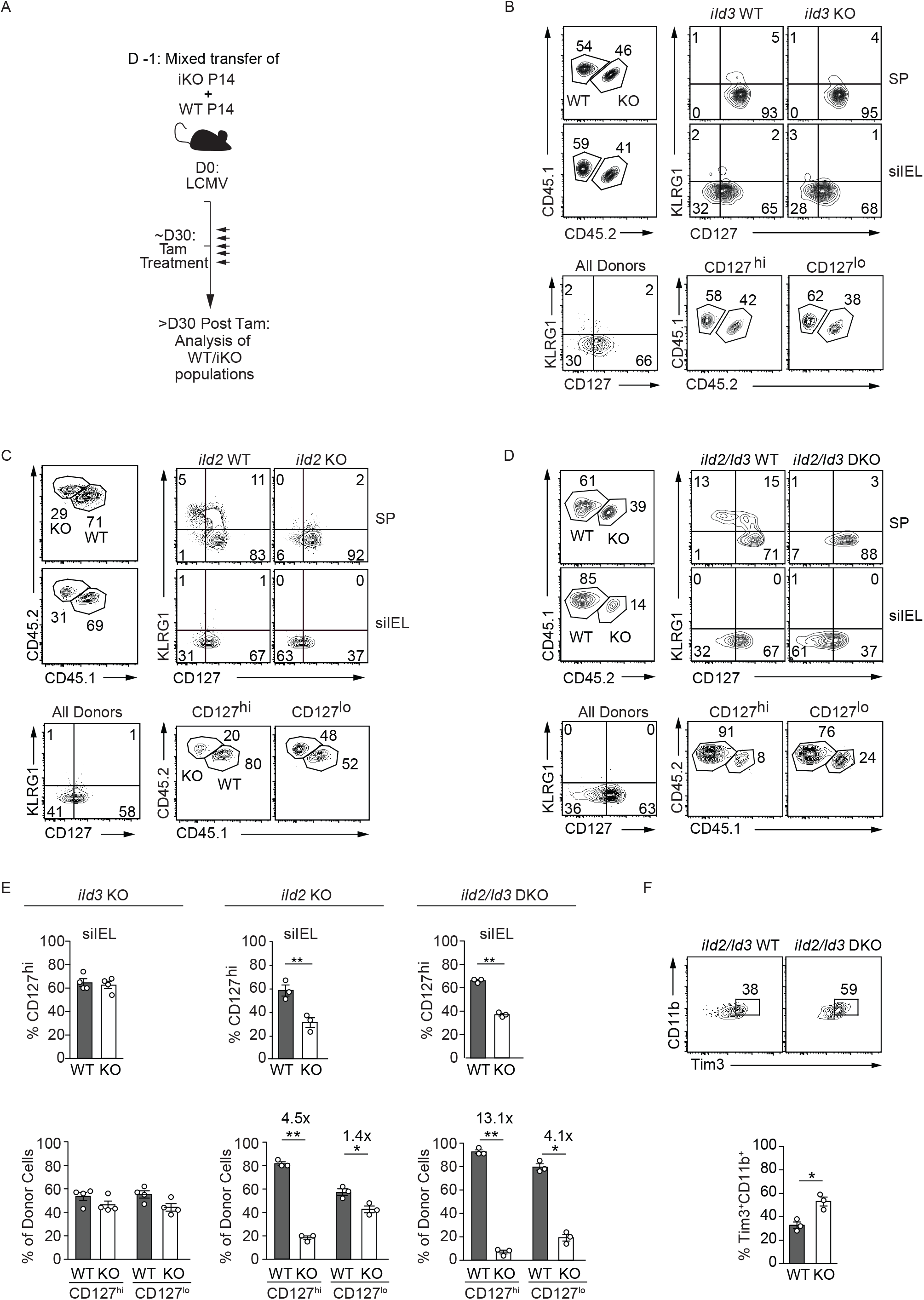
Id2 and Id3 mediate the maintenance of the long-lived siIEL CD8^+^ T cell population. **(A)** Schematic of experimental set-up. A mix of congenically distinct inducible knockout (iKO; Id2^f/f^-ERCre^+^, Id3^f/f^-ERCre^+^, Id2^f/f^Id3^f/f^ -ERCre^+^) and wild type (WT; ER-Cre^−^) P14 CD8^+^ T cells were transferred to wild-type recipients that were subsequently infected with LCMV. More than 30 days after infection, host mice were treated for 5 consecutive days with tamoxifen (Tam) to induce **(B)** *Id3*, **(C)** *Id2* or **(D)** *Id2* and *Id3* deletion. Transferred P14 CD8^+^ T cells from host spleen and siIEL were analyzed by flow cytometry >30 days after the last Tam treatment. Frequency of WT and iKO cells among P14 CD8^+^ T cells (top left) and corresponding KLRG1 and CD127 expression (top right) is represented. The proportion of WT and iKO P14 CD8^+^ T cells within the CD127^hi^ and CD127^lo^ populations are also shown (bottom). **(E)** Quantification of indicated populations is displayed. **(F)** Phenotype of *iId2/Id3* WT and DKO siIEL CD8^+^ T cells. Data are expressed as mean ± SEM. Numbers in plots represent frequency of cells in the indicated gate. All data are from one representative experiment of 2-3 independent experiments with n=3-5. Graphs show mean ± SEM; *p<0.05, **p<0.01.

### Id3^hi^ tumor-specific CD8^+^ T cells exhibit characteristics of Trm and progenitor-exhausted cells

It has previously been demonstrated that tumor infiltrating CD8^+^ lymphocytes (TIL) exhibit characteristics of Trm in some settings (Ganesan et al., 2017; Milner et al., 2017; Savas et al., 2018). Here, we utilized bulk- and sc-RNA-seq analysis to expand on this finding and investigate if tumor-specific CD8^+^ T cells exhibited similar heterogeneity to that observed in the siIEL CD8^+^ T cell population. P14 CD8^+^ T cells were isolated from the spleen and tumor of B16-GP_33-41_ melanoma-bearing mice 7 days after adoptive transfer and were processed for scRNA-sequencing (Figure 6A). Data from both the spleen and tumor were merged for VISION analysis. Despite heterogeneity within the TIL population, we found that TIL exhibited extensive enrichment of a tissue-residency gene-expression signature relative to tumor-specific T cells recovered from spleen (Figure 6A-B). We next utilized our Id3-GFP/Blimp1-YFP double-reporter system to evaluate if tumor-residing P14 CD8^+^ T cells can be subsetted analogously to siIEL tissue-resident cells. Indeed, P14 TIL consisted of Id3^lo^Blimp1^hi^, Id3^lo^Blimp1^lo^ and Id3^hi^Blimp1^lo^ populations, and RNAseq analysis revealed that Blimp1^hi^ TIL displayed a gene-expression pattern enriched for the Blimp1^hi^ siIEL CD8^+^ T cell signature, whereas Id3^hi^ TIL shared a gene-expression pattern with Id3^hi^ siIEL CD8^+^ T cells (Figure 6C). We extended this analysis by evaluating the relative enrichment of Blimp1^hi^ and Id3^hi^ Trm gene-expression signatures within individually profiled cells using VISION, and limited our analysis to only TIL (Figure 6D-E). Discrete patterns of enrichment were observed among TIL, indicating certain tumor-specific T cells exhibited a Blimp1^hi^ tissue-resident transcriptional profile while others had a more long-lived Id3^hi^ tissue-resident transcriptional profile.

**Figure 6.**
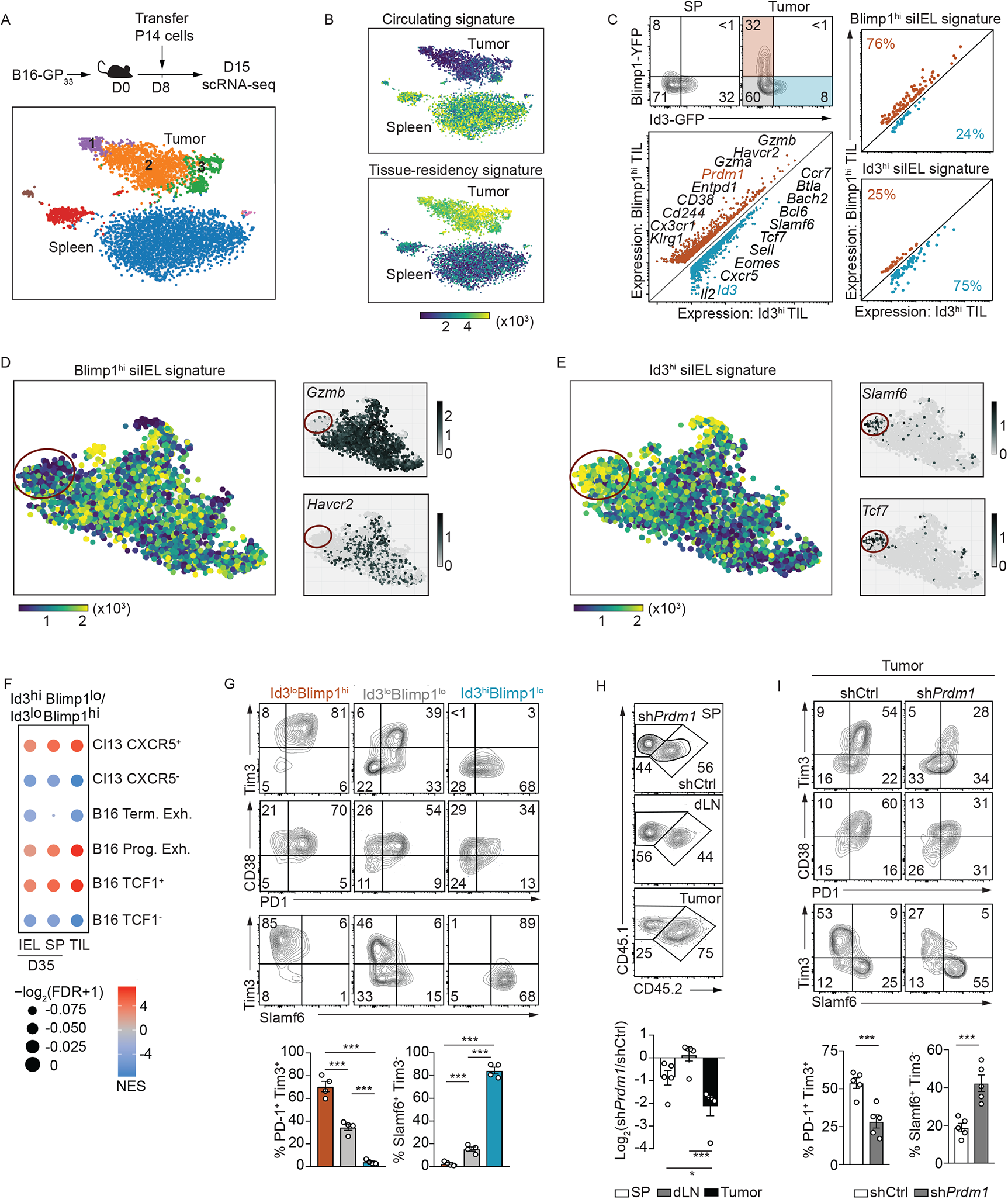
Blimp1 and Id3 expression distinguish distinct CD8^+^ T cell subsets in tumors. **(A)** Congenically distinct P14 CD8^+^ T cells were transferred into tumor-bearing mice, and seven days post adoptive transfer, P14 CD8^+^ T cells from spleens and tumors were sorted for scRNA-seq. VISION plot of P14 CD8^+^ T cells from tumors and spleens generated through VISION pipeline (DeTomaso et al., 2018). **(A)** VISION plot indicating relative enrichment of a core circulating signature (top) or enrichment of a core tissue-residency signature (bottom). (**C**) Representative flow cytometry plot demonstrating expression levels of Id3-GFP and Blimp1-YFP in P14 CD8^+^ T cells isolated from spleen or tumor (top). Expression plot from RNA-seq analysis of Blimp1^hi^Id3^lo^ and Blimp1^lo^Id3^hi^ P14 CD8^+^ T cells from tumors (bottom left). Relative expression levels of genes from Blimp1^hi^ and Id3^hi^ siIEL CD8^+^ T cells (from Figure 3B). (**D-E**) VISION plots indicating relative enrichment of the day 35 Blimp1^hi^ **(D)** or Id3^hi^ **(E)** siIEL CD8^+^ T cell signature (left) as well as relative expression of highlighted genes (right). Red circle indicates distinct cluster of TIL enriched with the Id3^hi^ Trm signature that also express elevated levels of genes upregulated in progenitor exhausted cells (Miller et al., 2019). **(F)** The CXCR5 (Im et al., 2016), B16 terminally/progenitor exhausted (Miller et al., 2019), and B16 TCF1 (Siddiqui et al., 2019) gene lists were used for gene set enrichment analyses in Id3^hi^ cells relative to Blimp1^hi^ cells. **(G)** Phenotype of Blimp1^lo^Id3^hi^, Blimp1^lo^Id3^lo^, and Blimp1^hi^Id3^lo^ TIL. **(H)** Congenically distinct P14 CD8^+^ T cells were transduced with a *Prdm1* shRNA encoding retrovirus (CD45.1^+^ P14 cells) or control shRNA (sh*Cd19*) encoding retrovirus (CD45.1^+^CD45.2^+^ P14 cells), mixed 1:1, and transferred into tumor bearing mice. Representative flow cytometry plots (top) and quantification (bottom) of the frequency of donor cells in the spleen, tumor draining lymph node, and tumor. Phenotype of *Prdm1*-deficient and control P14 CD8^+^ T cells from (H). Graphs show mean ± SEM of n=3-5 mice, from one representative experiment of 2 independent experiments. *p<0.05, ***p<0.005.

We noted that TIL enriched for the Blimp1^hi^ signature expressed elevated levels of *Havcr2* (encoding Tim3) and *GzmB*, whereas TIL enriched for the Id3^hi^ signature expressed elevated levels of *Slamf6* and *Tcf7*. It has previously been demonstrated that the exhausted CD8^+^ T cell population in the context of tumor or chronic infection are comprised of stem-like, progenitor exhausted (TCF1^hi^Slamf6^hi^ Cxcr5^hi^GzB^lo^Tim3^lo^CD38^lo^PD-1^mid^) and terminally exhausted (TCF1^lo^Slamf6^lo^Cxcr5^lo^GzB^hi^Tim3^hi^CD38^hi^ PD-1^hi^) subsets (Miller et al., 2019). Progenitor exhausted cells are reprogrammable and responsive to immunotherapies, whereas terminally exhausted cells primarily exist in a fixed state, less amenable to immunotherapy (Im et al., 2016; Miller et al., 2019; Siddiqui et al., 2019). As our scRNA-seq analysis identified a cluster of TIL enriched for the Id3^hi^ siIEL CD8^+^ T cell gene-signature as well as characteristic progenitor exhausted T cell markers, we performed GSEA to place the Id3^lo^Blimp1^hi^ and Id3^hi^Blimp1^lo^ CD8^+^ TIL subsets we identified within the context of already described populations (Im et al., 2016; Miller et al., 2019; Siddiqui et al., 2019). Indeed, Id3^hi^ TIL exhibited enrichment of gene-expression signatures associated with TCF1^+^ and progenitor exhausted T cell subsets identified in B16 tumors and CXCR5^+^ T cells responding to chronic LCMV infection, while Blimp1^hi^ TIL were most similar to TCF1^−^ and terminally exhausted TIL and CXCR5^−^ T cells of a chronic infection (Figure 6F). Flow cytometry analysis of TIL confirmed our gene-expression observations showing that the Id3^hi^ TIL population expressed higher levels of Slamf6 and lower amounts of GzmB and Tim3 compared to the corresponding Blimp1^hi^ subset (Figure 6G). Last, the Id3^hi^ TIL subset expressed higher levels of CD69 relative to Blimp1^lo^Id3^lo^ or Blimp1^hi^ subsets, consistent with a Trm phenotype (Figure S4).

Blimp1 is required for the formation of circulating TE and Tem (Kallies et al., 2009; Rutishauser et al., 2009) populations, supports Trm differentiation (Mackay et al., 2016), and regulates PD-1 expression during LCMV Cl13 infection (Shin et al., 2009). As we found that Blimp1 is required for optimal formation of the short-lived siIEL CD8^+^ T cell population following LCMV infection (Figure S3), we tested if Blimp1 regulates the accumulation and differentiation of TIL. We used a mixed transfer system wherein congenically distinct P14 CD8^+^ T cells were transduced with a retroviral vector encoding shRNAs targeting *Prdm1* or a negative control (*Cd19*), mixed at a 1:1 ratio and transferred into tumor bearing mice. One week following adoptive transfer, we found that the frequency of *Prdm1*-knockdown CD8^+^ T cells within the tumor was 4-fold lower than control CD8^+^ T cells. In contrast *Prdm1*-knockdown did not dramatically impact the accumulation of T cells within the spleen and lymph node (Figure 6H), thus highlighting a specific role for Blimp1 in controlling TIL accumulation (analogous to the role of Blimp1 in regulating siIEL Trm formation as in Figure S3). Further, Blimp1 was crucial for instructing differentiation of the terminally exhausted CD8^+^ T cell population within the tumor microenvironment as *Prdm1*-knockdown reduced the frequency of PD-1^+^Tim3^+^ CD8^+^ T cells but increased the proportion of a Slamf6^+^Tim3^−^ TIL (Figure 6I). These findings emphasize the transcriptional relationship between tissue-resident and tumor-resident CD8^+^ T cells and provide insight for manipulating the permanence and differentiation of TIL.

## Discussion

Paradigmatic studies have revealed that Trm are distinct from circulating memory CD8^+^ T cell populations and not conflated with the Tem population (Gebhardt et al., 2009; Mackay et al., 2013; Masopust et al., 2010; Wakim et al., 2012). This finding has shaped our understanding of the signals controlling Trm formation and function, yet a number of critical questions regarding the origin, fate, heterogeneity, and plasticity of Trm populations remain unanswered. Further, as the clinical relevance of Trm continues to widen from acute viral infections to settings of chronic inflammation and malignancy, clarification of the range of phenotypic and functional states exhibited by CD8^+^ T cells that reside in non-lymphoid tissues will provide a framework for understanding their regulation and identity in diverse contexts. While the single-cell analysis presented here and in the companion study by Kurd *et al*. (submitted) demonstrate that siIEL CD8^+^ T cells are transcriptionally distinct from circulating populations throughout infection, both studies also recognize considerable diversity in siIEL CD8^+^ T cell populations within a single timepoint as well. Further, our *in vivo* analyses confirmed clear inter-and intra-temporal heterogeneity within the siIEL CD8^+^ T cell population with a transcriptionally distinct, shorter-lived Blimp1^hi^ population most prominent early following both acute viral and bacterial infections as well as a longer-lived Id3^hi^ subset that emerged as the principal population at memory timepoints.

Functional heterogeneity within the circulating CD8^+^ T cell compartment affords adaptability to diverse pathogens (Chen et al., 2018). Comparable diversity within the tissue-resident T cell population is less appreciated, and the key molecules often used to define residency, CD69 and CD103, are almost uniformly expressed by siIEL Trm at memory timepoints (Casey et al., 2012; Mackay et al., 2013). Here, we define functional heterogeneity within the siIEL CD8^+^ T cell compartment, analogous to that described for the circulating CD8^+^ T cell pool. Our data indicate that the shorter-lived Blimp1^hi^KLRG1^int/lo^CD127^lo^ siIEL CD8^+^ T cell population ostensibly represent tissue-resident effector cells (TRE, during the effector phase of infection) or tissue-resident effector memory cells (Trem, during the memory phase of infection) as they are not only enriched for transcriptional signatures associated with more effector-like T cell populations (TE/Tem CD8^+^ T cells and TH1 CD4^+^ T cells) but also express elevated levels of effector molecule granzyme B. In contrast, the Id3^hi^KLRG1^lo^CD127^hi^ siIEL CD8^+^ T cells appear to be tissue-resident memory precursors (TrmP, during the effector phase of infection) or long-lived tissue-resident memory cells (Trm, during the memory phase of infection) as they share transcriptional signatures with other memory or memory-like populations (MP/Tcm CD8^+^ T cells and Tfh CD4^+^ T cells) and have increased memory potential and multipotency, with the capacity to generate circulating and Trm populations following re-infection. Trm can rapidly proliferate *in situ* following re-infection (Beura et al., 2018a; Park et al., 2018), and in certain contexts, may exit the tissue and rejoin the circulating memory pool (Beura et al., 2018b; Masopust et al., 2006). We propose that it is the more stem-like Id3^hi^ Trm subset that undergoes local proliferation during secondary infection and differentiates into the circulating ex-Trm population.

A number of reports have previously inferred heterogeneity exists within the tissue-resident compartment. Following *Yersinia pseudotuberculosis* infection, both CD103^−^ and CD103^+^ Trm populations can be found within the lamina propria that have been suggested to contribute to immediate pathogen clearance or participate in the secondary response (Bergsbaken and Bevan, 2015; Bergsbaken et al., 2017), consistent with our results highlighting a tissue-resident population with effector-like qualities (TRE/Trem). Further supporting the idea of tissue-resident T cells with distinct functions, diverse subsets of human Trm were recently identified in multiple tissues (Kumar et al., 2018). These subsets of human Trm include a population of cells with features of longevity and quiescence that demonstrate a heightened proliferative capacity, as well as a Trm population with elevated effector capacity, analogous to the Id3^hi^ and Blimp1^hi^ populations we report here (Kumar et al., 2018).

Drawing comparisons from the circulating CD8^+^ T cell populations, we examined the requirement of previously defined T cell fate-specifying transcriptional regulators for differentiation and maintenance of the siIEL CD8^+^ T cell populations. We found that pro-effector transcriptional regulators (Blimp1, Id2, and T-bet) direct the differentiation of siIEL CD8^+^ T cells, consistent with previous studies describing that Blimp1 and T-bet control Trm formation (Laidlaw et al., 2014; Mackay et al., 2016; Mackay et al., 2015). However, here, we expand on these findings in the context of the newly defined siIEL CD8^+^ T cell heterogeneity, and demonstrate that loss of Blimp1, Id2 or T-bet impairs formation of Blimp1^hi^KLRG1^hi^ TRE while the differentiation of the Id3^hi^KLRG1^lo^ precursors of Trm were less affected or even enhanced (Laidlaw et al., 2014; Mackay et al., 2015). Notably, both Id2 and Id3 were found to enforce siIEL CD8^+^ T cell homeostasis, with the CD127^hi^ longer-lived Trm subset showing a greater dependency on these transcriptional regulators than CD127^lo^ Trem. At day 60 of infection, Id2 is upregulated in the long-lived Id3^hi^ siIEL population, and thus the role of Id2 shifts from supporting TRE formation to maintaining the long-lived Trm population. In the circulation, Id3 maintains long-lived CD8^+^ T cells (Ji et al., 2011; Yang et al., 2011) and is repressed by Blimp1 (Ji et al., 2011), yet Id2 regulation sustains the more effector-like memory CD8^+^ T cell populations (Omilusik et al., 2018). Thus, while circulating and resident populations share some underlying transcriptional dependencies, the siIEL CD8^+^ T cell compartment is also distinctly dependent on unique transcriptional signals as well. Consistent with our finding that divergent transcriptional programs direct short- and long-lived siIEL CD8^+^ T cell subsets, a recent study described inverse expression of Blimp1 and Hobit in skin Trm and suggested early effector populations rely on Blimp1 regulation but memory-like populations require Hobit (Mackay et al., 2016). Conceptualization of Trm heterogeneity in comparison to the well-defined circulating memory T cell subsets provides context for understanding their functional differences. Kurd *et al*. (submitted) define gene modules that are distinct between siIEL and splenic CD8^+^ T cells and identify differentially expressed genes between distinct cell clusters within individual time points (day 4, 60 and 90) of LCMV infection beyond those described here that include transcription factors, chemokines, cytokines and associated signaling molecules, cell survival molecules, and costimulatory receptors, indicating substantially more complex heterogeneity. Thus, while parallels can be drawn with circulating memory subsets, it is clear that the siIEL CD8^+^ T cell population exhibits unique tissue-specific features and includes cells with a spectrum of function and memory potential.

Trm are present in highly diverse tissue environments so it is not surprising that tissue-specific variability also exists. Importantly, while Blimp1-expressing Trm were found in all tissues that we examined, Id3^hi^ Trm were present within the gut and skin suggesting that these barrier tissues confer a tissue milieu that promotes and maintains Id3 expression. Trm cells within epithelial compartments require TGFβ for optimal differentiation and maintenance as it induces upregulation of CD103 to facilitate tissue retention (Casey et al., 2012; Mackay et al., 2013; Zhang and Bevan, 2013). TGFβ has also been shown to induce Id3 expression in various lymphoid populations (Kee et al., 2001), and thus may contribute to Id3 expression within the gut and skin Trm as well. This suggests an additional layer of Trm heterogeneity exists and that transcriptional programs within Trm are induced and supported by tissue-specific cues allowing Trm to adapt to the conditions posed by the local environment.

Accumulating evidence indicates that certain populations of TIL exhibit characteristics of tissue-resident cells, and the presence of Trm-like TIL is linked to positive prognoses in multiple malignancies and likely confers durable anti-tumor immunity (Ganesan et al., 2017; Miller et al., 2019; Savas et al., 2018). Here, we provide evidence that despite heterogeneity, nearly all tumor-specific T cells are enriched for a core tissue-residency gene-expression signature. Further, we find that sub-populations of TIL transcriptionally and phenotypically resemble Blimp1^hi^ Trem and Id3^hi^ Trm. Consistent with Id3 marking multipotent or memory-like siIEL CD8^+^ T cells, Id3^hi^ Trm-like cells in tumors exhibit features of the recently described progenitor exhausted cells, a multipotent population of TIL that are responsive to immunotherapies such as cancer vaccination and immune checkpoint blockade (Im et al., 2016; Miller et al., 2019; Siddiqui et al., 2019). Exploring strategies to bolster this Id3^hi^ Trm-like subset within tumors will likely be important to increase efficacy of these treatments. Our findings provide an additional layer in the understanding of the diverse subsets of T cells residing in the tumor microenvironment, which may help facilitate manipulation of T cell activity to enhance cancer immunotherapy efficacy. Taken together, both circulating and resident CD8^+^ T cells in the context of infection and cancer display phenotypic and functional heterogeneity, ranging from multipotent to terminally differentiated, endowing the immune system with flexible protection at sites of imminent pathogen exposure or ongoing disease.

## Materials and methods

### Mice, adoptive cell transfer, infections, tumor model, skin inflammation, and tamoxifen treatment

All mouse strains were bred and housed in specific pathogen–free conditions in accordance with the Institutional Animal Care and Use Guidelines of the University of California San Diego. Blimp1-YFP mice (stock #008828;The Jackson Laboratory), Id3-GFP mice (Miyazaki et al., 2011), Id2-YFP mice (Yang et al., 2011), *Prdm1*^fI/fl^ Granzyme B-Cre mice (Rutishauser et al., 2009), *Tbx21*^+/−^ mice (stock #004648, The Jackson Laboratory), *Id2*^fl/fl^ mice (Niola et al., 2012), *Id3*^fl/fl^ mice (Guo et al., 2011), CD4-Cre mice (stock #017336; The Jackson Laboratory), *Rosa26*Cre-ERT2 (ERCre) (Hess Michelini et al., 2013), P14 mice (with transgenic expression of H-2D^b^-restricted TCR specific for LCMV glycoprotein gp_33-41_), OT-I mice (with transgenic expression of H-2K^b^-restricted TCR specific for ovalbumin peptide 257-264; stock #003831; The Jackson Laboratory), CD45.1^+^ and CD45.1.2^+^ congenic mice were bred in house. Both male and female mice were used throughout the study, with sex and age matched T cell donors and recipients (or female donor cells transferred into male recipients).

Wild-type, Blimp1-YFP, Id3-GFP, Id2-YFP/Id3-GFP or Blimp1-YFP/Id3-GFP P14 CD8^+^ T cells congenically distinct for CD45 were adoptively transferred at 5×10^4^ cells per recipient mouse. For cotransfers, *Prdm1*^fl/fl^-granzyme B-Cre, *Id2*^fl/fl^ CD4-Cre, *Id2*^fl/fl^ ER-Cre, *Id3*^fl/fl^ ER-Cre or I*d2*^fl/fl^*Id3*^fl/fl^ ER-Cre and corresponding control P14 CD8^+^ T cells were mixed in a 1:1 ratio and adoptively transferred at 5×10^4^ total cells per recipient mouse. Alternatively, *Tbx21*^+/−^ and *Tbx21*^+/+^ OT-I CD8^+^ T cells were mixed and similarly transferred to congenically distinct recipients. Mice were then infected with 2×10^5^ pfu lymphocytic choriomeningitis virus-Armstrong (LCMV) or LCMV-Armstrong expressing ovalbumin (LCMV-OVA) by intraperitoneal injection or with 1×10^10^ *Listeria monocytogenes* expressing GP_33_ (LM-GP_33_) by oral gavage. For skin Trm studies, splenocytes from Id3-GFP P14 mice were *in vitro* activated with 10 nM gp33 peptide and expanded in 100U/mL IL-2 for 4 days, then 5×10^6^ P14 cells were transferred intravenously to congenically distinct recipients that were treated the same day with 15μl of 0.3% 1-fluoro-2,4-dinitrobenzene (DNFB) in acetone/oil (4:1) applied to the flank skin (Davies et al., 2017). For ER-Cre-mediated deletion of floxed alleles, 1 mg tamoxifen (Cayman Chemical Company) emulsified in 100 μl of sunflower seed oil (Sigma-Aldrich) was administered by intraperitoneal injection for 5 consecutive days after 30 days of infection.

B16-GP_33-41_ cells (5 x10^5^) were transplanted subcutaneously into the right flank of wild-type mice. After tumors became palpable, 7–8 days after transplantation, 1-2×10^6^ *in vitro* expanded Blimp1-YFP/Id3-GFP P14 CD8^+^ T cells,WT P14 cells (for scRNAseq), or retrovirally transduced cells were transferred intravenously. For *Prdm1* RNAi studies, congenically distinct sh*Prdm1* and shCtrl P14 cells were mixed 1:1 prior to transfer into tumor-bearing mice. Tumors were monitored daily and mice with ulcerated tumors or tumors exceeding 1500 mm^3^ in size were euthanized, in accordance with UCSD IACUC. Tumor infiltrating lymphocytes (TIL) were isolated as previously described (Milner et al.) one week following adoptive transfer.

### Preparation of single cell suspensions

Single-cell suspensions were prepared from spleen or lymph node by mechanical disruption. For small intestine preparations, Peyer’s patches were excised, luminal contents were removed, and tissue was cut longitudinally then into 1cm pieces. The gut pieces were incubated while shaking in 10% HBSS/HEPES bicarbonate solution containing 15.4mg/100μL of dithioerthritol (EMD Millipore) at 37°C for 30 minutes to extract siIEL. For lamina propria lymphocyte isolation, gut pieces were further treated with 100U/ml type I collagenase (Worthington Biochemical) in RPMI-1640 containing 5% bovine growth serum, 2 mM MgCl_2_ and 2 mM CaCl_2_ at 37°C for 45 minutes. Brain, white adipose tissue, salivary gland, kidney, and tumor were cut with scissors into fine pieces then incubated while shaking with 100U/ml type I collagenase as above. Epidermal cells were isolated by floating skin dermis side down on 0.3% trypsin in 150mM NaCl, 6mM KCl, 6mM glucose, pH7.6 for 2hr at 37°C. The epidermis was then separated from the dermis and incubated with shaking for a further 10min at 37°C in 0.3% trypsin solution. Cells were then filtered through a sera separa column to remove any remaining tissue pieces. Lymphocytes from all tissue but skin, spleen and lymph node were purified on a 44%/67% Percoll density gradient.

### Antibodies, flow cytometry, and cell sorting

The following antibodies were used for surface staining (all from eBioscience unless otherwise specified): CD5 (53-7.3, BD Pharmingen), CD8 (53-6.7), CD11b (M1/70) CD27 (LG-7F9), CD45.1 (A20-1.7), CD45.2 (104), CD69 (H1.2F3, Biolegend), CD103 (2E7), CD127 (A7R34), CD160 (CNX46-3), CX3CR1(SA011F11, Biolegend), KLRG1 (2F1), PD-1(J43), Slamf6 (13G3), Tim3 (RMT3-23). Cells were incubated for 30 min at 4°C in PBS supplemented with 2% bovine growth serum and 0.1% sodium azide. Intracellular staining was performed using the BD Cytofix/Ctyoperm Solution Kit (BD Biosciences) and the following antibodies (all from eBioscience unless otherwise specified): Eomes (Dan11mag), Granzyme B (GB12, Invitrogen), IFN-*γ* (XMG1.2), IL-2 (JES6-5H4), T-bet (4B10), TCF1 (C63D9, Cell Signaling), TNF*α* (MP6-XT22). For cytokine staining, siIEL CD8^+^ T cells were incubated for 3 hours at 37°C in RPMI-1640 media containing 10% (v/v) bovine growth serum with 10 nM gp_33-41_ peptide and Protein Transport Inhibitor (eBioscience). CD107a (1D4B, BD Pharmingen) antibody was included in the media for the entirety of the stimulation to detect surface expression as a surrogate of degranulation. Stained cells were analyzed using LSRFortessa or LSRFortessa X-20 (BD) and FlowJo software (TreeStar). All sorting was performed on BD FACSAria or BD FACSAria Fusion instruments.

### Bulk RNA-Sequencing

On day 55 of LCMV infection,1×10^3^ P14 cells from the spleen (CD62L^+^ Tcm and CD62L^−^ Tem) or CD62L^−^ CD103^+^ P14 cells from the siIEL were sorted into TCL buffer (Qiagen) with 1% 2-Mercaptoethanol. On day 7 or 35 day of LCMV infection, 1×10^3^ Blimp1^lo^Id3^hi^, Blimp1^hi^Id3^lo^ Blimp1^lo^Id3^lo^ P14 cells from the spleen, mesenteric lymph node or small intestine siIEL were sorted similarly. For tumor studies, 1×10^3^ Blimp1^lo^Id3^hi^ and Blimp1^hi^Id3^lo^ P14 cells were sorted from B16-GP_33_ tumors or spleens 7 days post adoptive transfer (14 days post-transplant). For all samples, polyA^+^ RNA was isolated and RNA-seq library preparation as well as RNA-seq analysis were carried out as described in (https://www.immgen.org/Protocols/11Cells.pdf). The heatmaps in Figure 1a were generated using Morpheus (https://software.broadinstitute.org/morpheus); 3490 differentially expressed (D.E.) transcripts (>2-fold, expression threshold >10) between all three populations of Tcm, Tem and Trm were identified through the Multiplot Studio module within Genepattern. The D.E. genes were then ordered through K-means clustering (set to 6 clusters) with the following modifications within Morpheus: metric=one minus pearson correlation; maximum number of iterations=1000. For the transcription factor heatmap in Figure 1A (right), transcription factors were cherry picked from the larger heatmap (left) based on known or predicted roles in regulating Tcm or Tem fate. RNA-sequencing was performed on duplicate samples as such: day 55 LCMV sorts, spleens or siIEL cells from two mice were pooled for one replicate; for day 8 LCMV and day 35 LCMV sorts, 2-5 spleens, mLNs or siIEL cells were pooled per one replicate; for TIL, spleens and tumors from two mice were pooled per replicate.

For triwise plots, 2-fold differentially expressed genes (between D55 Tcm, Tem and Trm; expression threshold >10) were filtered in Multiplot Studio based on the following gene lists: ‘Effector,’ ‘Memory,’ ‘Core Residency Signature,’ and ‘Core Circulating Signature.’ Expression values from filtered genes were then log_2_ transformed. Subsequently, triwise plots and rose plots were generated with the log_2_ transformed data using the Triwise R package (van de Laar et al., 2016) and following https://zouter.github.io/triwise/rd.html. The ‘Core Residency Signature’ and ‘Core Circulating Signature’ are from (Milner et al., 2017). The ‘effector’ and ‘memory’ signatures were generated using the ImmGen database: https://www.immgen.org/. Specifically, utilizing default settings (expression threshold >120, 2-fold change) in the Population Comparison module, we identified differentially expressed genes between T_8Eff_Sp_OT1_d8_LisOva and T_8Mem_Sp_OT1_d100_LisOva. Genes upregulated in T_8Eff_Sp_OT1_d8_LisOva comprised the ‘effector’ signature and genes upregulated in T_8Mem_Sp_OT1_d100_LisOva comprised the ‘memory’ signature.

RNAseq expression plots were generated in Multiplot Studio within GenePattern (expression threshold >10, 1.5-fold differentially expressed). In Figure 6C, the Id3^hi^ siIEL signature is comprised of genes upregulated 1.5-fold in both D7 Id3^hi^ IEL vs. D7 Blimp1^hi^ IEL and D35 Id3^hi^ IEL vs D35 Blimp1^hi^ IEL comparisons, whereas the Blimp1^hi^ siIEL signature is comprised of genes upregulated 1.5-fold in both D7 Id3^hi^ IEL vs D7 Blimp^hi^ IEL and D35 Id3^hi^ IEL vs D35 Blimp1^hi^ IEL comparisons. For Figure 6D-E, the Blimp1^hi^ signature consists of genes upregulated in D35 Id3^hi^ IEL vs. D35 Blimp1^hi^ IEL, whereas the Blimp1^hi^ signature consists of genes downregulated in D35 Id3^hi^ IEL vs D35 Blimp1^hi^ IEL.

Gene set enrichment analysis was performed on each cell subset using the GSEA Preranked gene list tool with 1000 permutations and a weighted enrichment statistic. Gene sets were obtained by taking the top differentially expressed genes from microarray (GSE9650) or RNA-seq (GSE84105, GSE122713) data from the GEO database. Normalized enrichment scores (NES) and false discovery rate (FDR) were visualized using the ggplot2 package in R. For principal component analysis (PCA) plots, normalized counts were condensed to contain the top 535 differentially expressed genes between all samples with a padj < 0.05. PCA was performed using the prcomp function in R after centering and scaling the normalized counts.

### 10x Genomics library preparation and sequencing

Activated P14 T cells (CD8^+^V*α*2^+^CD45.1^+^CD44^+^) were sorted from the spleen or siIEL and resuspended in PBS+0.04% (w/v) bovine serum albumin. Approximately 10,000 cells per sample were loaded into Single Cell A chips (10x Genomics) and partitioned into Gel Bead In-Emulsions (GEMs) in a Chromium Controller (10x Genomics). Single cell RNA libraries were prepared according to the 10x Genomics Chromium Single Cell 3’ Reagent Kits v2 User Guide and sequenced on a HiSeq4000 (Illumina).

#### Infection studies

##### Single-cell RNA-seq mapping

Reads from single-cell RNA-seq were aligned to mm10 and collapsed into unique molecular identifier (UMI) counts using the 10X Genomics Cell Ranger software (version 2.1.0). All samples had sufficient numbers of genes detected (>1000), a high percentage of reads mapped to the genome (>70%), and sufficient number of cells detected (>1000).

##### Cell and gene filtering

Raw cell-reads were then loaded to R using the cellrangerRkit package. The scRNA-seq dataset was then further filtered based on gene numbers and mitochondria gene counts to total counts ratio. Only cells with > 400 genes, UMI > 0, and 0.5% ~ 30% of their UMIs mapping to mitochondria genes were kept for downstream analysis. To ensure that memory requirements for all downstream analyses did not exceed 16Gb and that the samples with more cells would not dominate the downstream analysis, we randomly selected a portion of the cells that passed filtering for downstream analysis. We randomly selected 2000 cells from each library for downstream analysis. After cell filtering and sampling, we filtered genes by removing genes that did not express > 1 UMI in more than 1% of the total cells.

##### Single-cell RNA-seq dataset normalization and pre-processing

Five cell-gene matrices were generated:

1. Raw UMI matrix.
2. UPM matrix. The raw UMI matrix was normalized to get UMIs per million reads (UPM), and was then log2 transformed. All downstream differential analysis was based on the UPM matrix. The prediction models were also based on the UPM matrix, as other normalizations are very time-consuming for large datasets.
3. MAGIC matrix. UPM matrix was further permuted by MAGIC (van Dijk et al., 2018) R package Rmagic 1.0.0 was used, and all options were kept as default. MAGIC aims to correct the drop-out effect of single-cell RNA-seq data; thus, we used MAGIC-corrected matrix for visualizing the gene expression pattern rather than using the UPM matrix. All gene expression overlaid on TSNE plots were based on the MAGIC matrix.
4. Super cell matrix. We merged 50 cells to create a ‘super’ cell and used the super cell matrix as the input for cell type annotation analysis. This approach enabled us to bypass the issue of gene dropouts with scRNA-seq and make the data comparable to bulk RNA-seq. We first calculated the mutual nearest neighbor network with k set to 15, and then cells that were not mutual nearest neighbors with any other cells were removed as outliers. We randomly selected ‘n’ cells in the UPM matrix as the seed for super cells. The expression of each super cell was equal to the average expression of its seed and the 50 nearest neighbor cells of its seed. We derived 7400 super cells from the dataset, so each single cell was covered ~10 times.

##### Single-cell RNA-seq dataset dimension reduction

Top variable genes, PCA, and tSNE were calculated by Seurat version 2.3.4 functions: FindVariableGenes, RunPCA, and RunTSNE (Butler et al., 2018). Only the top 3000 genes were considered in the PCA calculation and only the top 25 principal components (PCs) were utilized in tSNE. Louvain clustering was performed by Seurat’s FindClusters function based on the top 25 PCs, with resolution set to 2. The scaled UPM matrix was used as input in the calculation.

##### Annotating single-cells with bulk RNA-seq signatures

The log_2_ TPM data from bulk RNA-seq datasets were compared with the scRNA-seq super cell matrix. Bulk cell population RNA-seq samples were first grouped into different sets according to their mutual similarities. For each bulk RNA-seq sample set, the mean expression was first calculated. The 1^st^ correlation was calculated between all the super cells and the mean expression from the bulk RNA-seq dataset. Based on the distribution of the 1^st^ correlation, we were able to identify a group of super cells that were most similar to the mean expression of the bulk sample. To further identify the small differences between bulk RNA-seq expression within a given set, we removed the set mean from the bulk RNA-seq and the mean from the most similar group of super cells, and then calculated the 2nd correlation between the super cells and bulk RNA-seq. Based on the 2nd correlation, we annotated the super cells with each bulk sample label.

##### Comparing gut and spleen CD8^+^ T cells

In order to compare the gut cells and spleen cells from the perspective of effector-memory differentiation, we performed a 3-step analysis. First, we calculated the differentially expressed gene between effector and memory T cells. We then performed principal component analysis on the gene expression of the DE genes of all spleen cells. Finally, we mapped both spleen cells and gut cells to the top two principle components calculated in step 2. Therefore, we can examine the heterogeneity of gut cells in a low-dimensional space that reveals the most significant difference between spleen effector and memory CD8 T cells.

##### Tumor studies

Analysis was performed similarly to the infection studies described above. Sequencing was performed on duplicate samples of splenocytes and TIL, wherein each replicate was pooled from 2-3 mice. Briefly, reads were aligned to mm10 and collapsed into UMI counts using the 10X Genomics Cell Ranger software. All samples had sufficient numbers of genes detected (>1000), a high percentage of reads mapped to the genome (>70%), and sufficient number of cells detected (>1000).

Raw cell-reads were then loaded to R using the Seurat package version 2.3.4 for further processing and quality control. The scRNA-seq dataset was filtered based on gene numbers and mitochondria gene counts to total counts ratio. Only cells with > 200 genes, UMI > 0, and 0%~5% of their UMIs mapping to mitochondria genes were kept for downstream analysis. Raw UMI matrix was normalized using Seurat function NormalizeData with the “LogNormalize” option, which normalized the feature expression measurements for each cell by the total expression, multiplies this by a scale factor (10,000 by default), and log-transforms the result. Normalized matrix was scaled using Seurat function ScaleData which regressed out cell-cell variation in gene expression driven by number of UMI and mitochondrial gene expression. VISION (DeTomaso et al., 2018) analysis was performed on the scaled and normalized Seurat object. VISION computes a K-nearest-neighbor graph and PCA dimensionality reduction for the Seurat object. The gene signatures used are described above in Bulk RNA-sequencing methods. For each signature, an overall score was calculated for every cell summarizing the expression of genes in the signature.

### Statistics

Statistical analysis was performed using GraphPad Prism software. Two-tailed paired or unpaired t-test was used for comparisons between groups. P values of <0.05 were considered significant.

### Data availability

The LCMV infection single-cell RNA sequencing data are available for download on the GEO data repository with accession number GSE131847. All bulk RNA-seq datasets and tumor single-cell RNA-seq will be uploaded to the GEO repository prior to publication.

### Code availability

Codes are available at https://github.com/Arthurhe/scRNA_CD8T_Goldrath_Project.

## Acknowledgements

This work was funded by the NIH grants AI132122 (A.W.G., G.W.Y., J.T.C.); K99 CA234430-01 (J.J.M.). Single-cell RNA-sequencing using the 10X Genomics platform was conducted at the IGM Genomics Center, University of California, San Diego, La Jolla, CA and supported by grants P30KC063491 and P30CA023100. We thank the Flow Cytometry Core at the La Jolla Institute for Allergy and Immunology for their assistance with cell sorting, and the Chang, and Goldrath laboratories for technical advice, helpful discussion, and critical reading of the manuscript.

---

**Figure S1.**
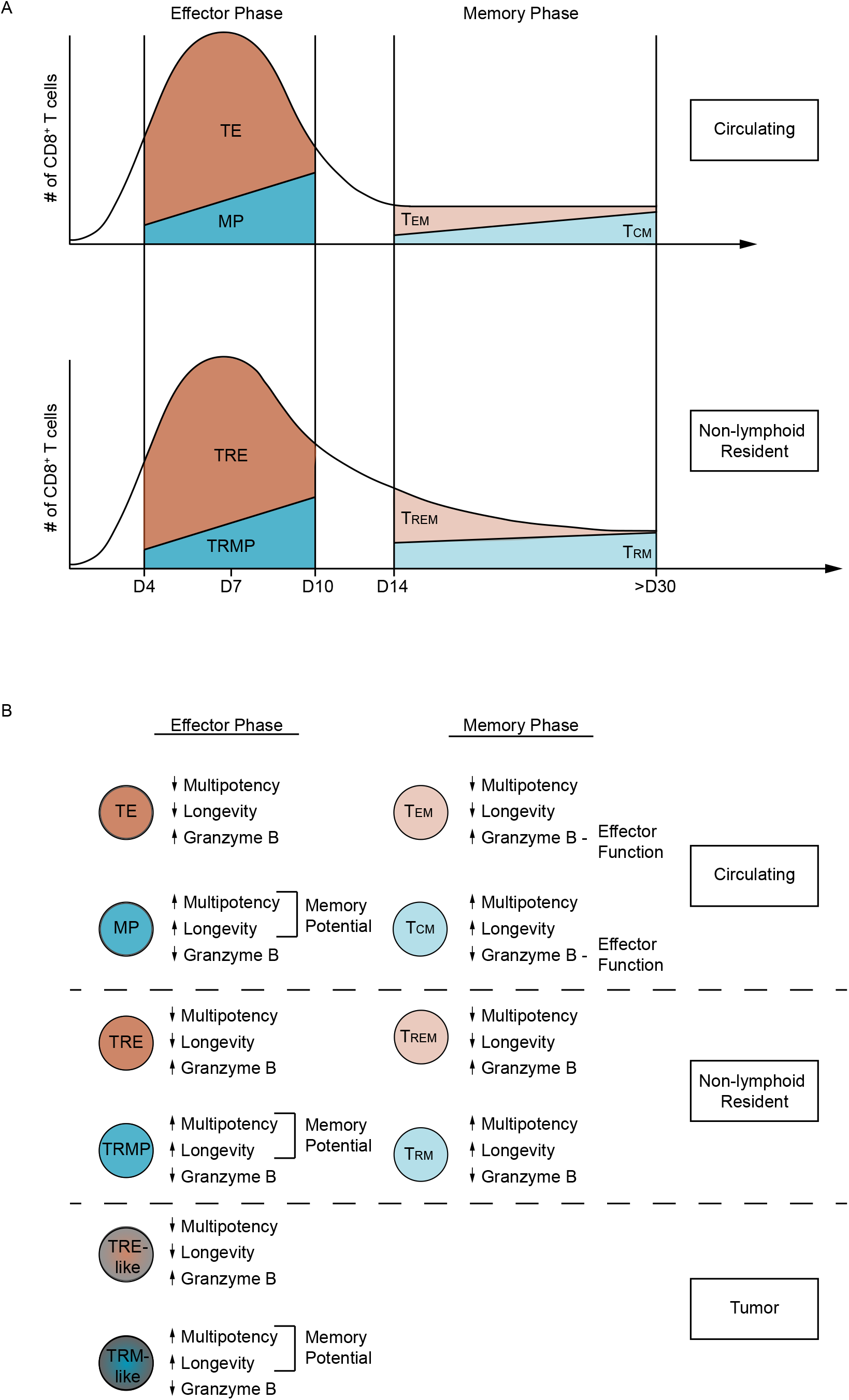
Model of heterogeneity in CD8^+^ T cell populations. **(A)** Non-lymphoid resident populations during effector and memory phases of infection exhibit heterogeneity analogous to circulating populations. **(B)** siIEL CD8^+^ T cells during early infection timepoints are enriched with qualities of effector cells whereas long-lived siIEL CD8^+^ T cells at subsequent timepoints are enriched with memory attributes. siIEL CD8^+^ T cells also exhibit intra-temporal heterogeneity at both effector and memory phases of infection with Blimp1 and Id3 distinguishing distinct tissue-resident effector or tissue-resident effector memory cells (TRE and Trme) as well as tissue-resident memory precursors and tissue-resident memory cells (TrmP and Trm). Trme- and Trm-like CD8^+^ T cell subsets can also be identified within the tumor microenvironment.

**Figure S2.**
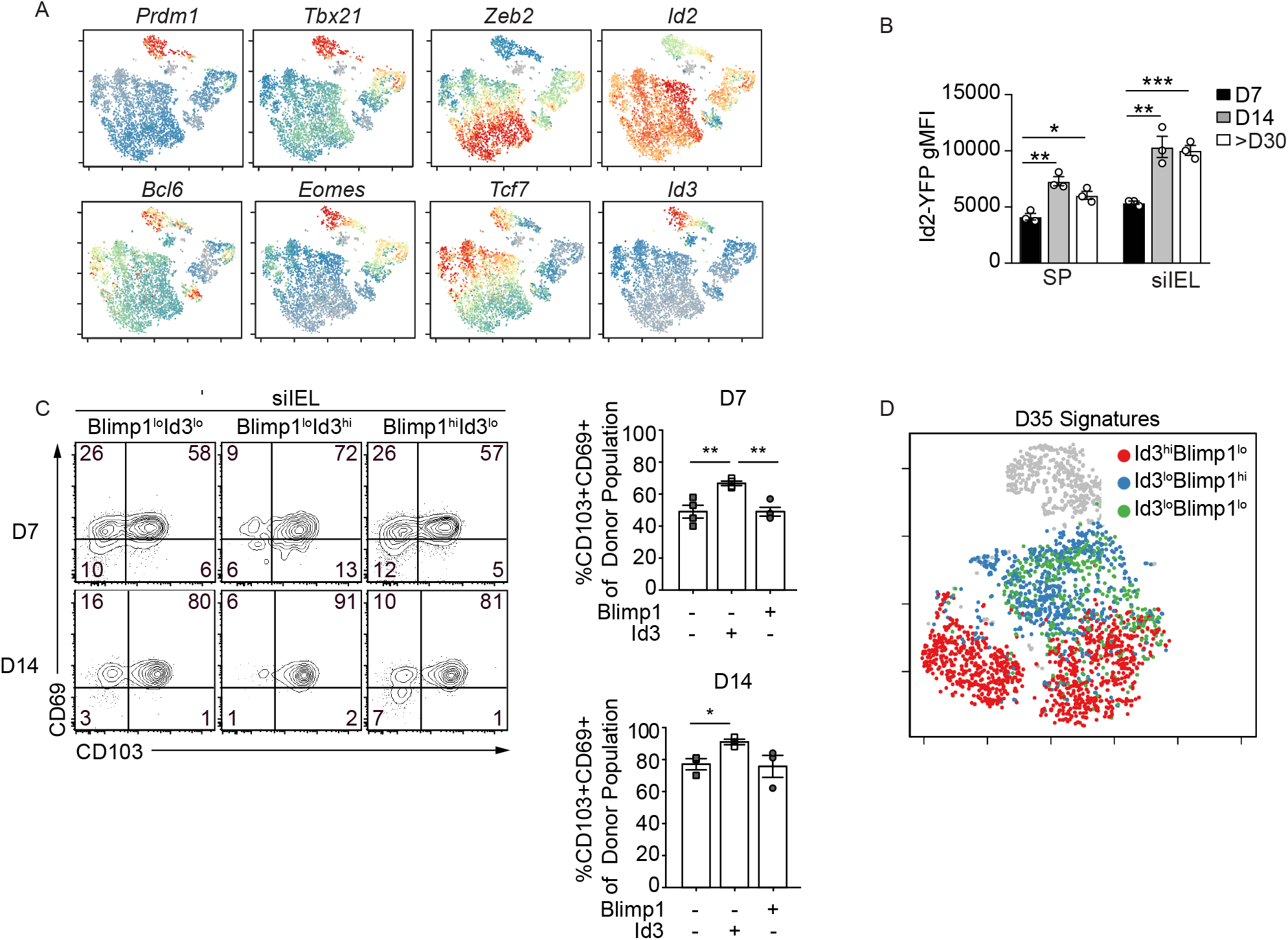
Heterogeneity in anti-viral CD8^+^ T cell populations. **(A)** bP14 CD8^+^ T cells were transferred into congenically distinct hosts that were subsequently infected with LCMV. Donor cells from the spleen and siIEL were sorted over the course of infection for scRNA-seq. tSNE plots of cells from the spleen over all infection timepoints colored by intensity of indicated transcriptional regulator**. (B)** Id2-YFP/Id3-GFP P14 CD8^+^ T cells were transferred into congenically distinct hosts that were infected with LCMV. At indicated times of infection, Id2-YFP reporter expression was analyzed in the P14 cells isolated from the spleen and siIEL by flow cytometry and the gMFI is quantified. **(C)** Congenically distinct Blimp1-YFP/Id3-GFP P14 CD8^+^ T cells were transferred to wild-type hosts that were subsequently infected with LCMV. CD103 and CD69 expression on indicated populations is shown. **(D)** As in A, tSNE plots of cells from the siIEL over all timepoints colored by intensity of indicated transcriptional signatures at day 35 following LCMV infection. Numbers in plots represent the frequency of cells in the indicated gate. All data are from one representative experiment of 2 independent experiments with n=3. Graphs show mean ± SEM; *p<0.05, **p<0.01, ***p<0.001.

**Figure S3.**
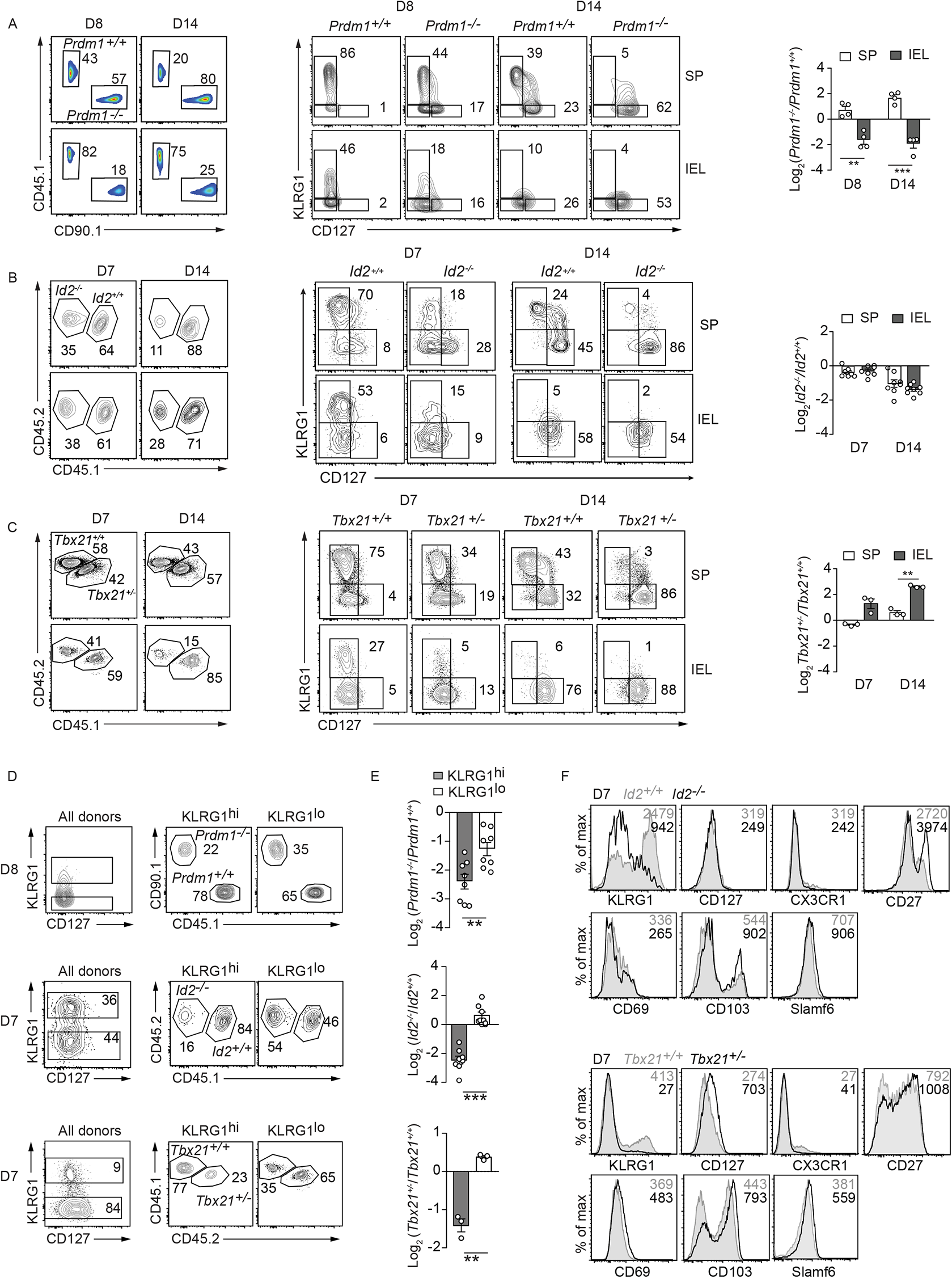
Pro-effector transcriptional regulators mediate siIEL CD8^+^ T cell formation. Mixed transfer of congenically distinct *Prdm1*^f/f^ GzmB-Cre^+^ (*Prdm1*^−/−^) and *Prdm1*^+/+^, *Id2^f/f^* CD4-Cre^+^ (*Id2^−/−^*) and Id2^f/f^ CD4-Cre^−^ (*Id2^+/+^*) or Tbx21^+/−^ and Tbx21^+/+^ P14 CD8^+^ T cells into recipient mice subsequently infected with LCMV. Donor P14 cells from host spleen and siIEL were analyzed by flow cytometry at indicated times of infection. **(A-C)** Frequency of indicated donor populations among CD8^+^ T cells and corresponding quantification is shown (left). KLRG1 and CD127 expression on indicated donor populations is represented (middle/right). **(D)** KLRG1^hi^ and KLRG1^lo^ populations of all donor P14 CD8^+^ T cells from the siIEL (left) and the frequency of the donor cells within these populations (right) is shown. **(E)** Quantification of populations from D. **(F)** Expression of indicated molecules is compared between *Id2*^−/−^ and *Id2*^+/+^ (top) or *Tbx21*^+/−^ and *Tbx21*^+/+^ (bottom). Numbers in plots represent the frequency (A-D) or gMFI (F) of cells in the indicated gate. All data are from one representative experiment of 2 independent experiments with n=3-5. Graphs show mean ± SEM; **p<0.01, ***p<0.001.

**Supplemental Figure 4.**
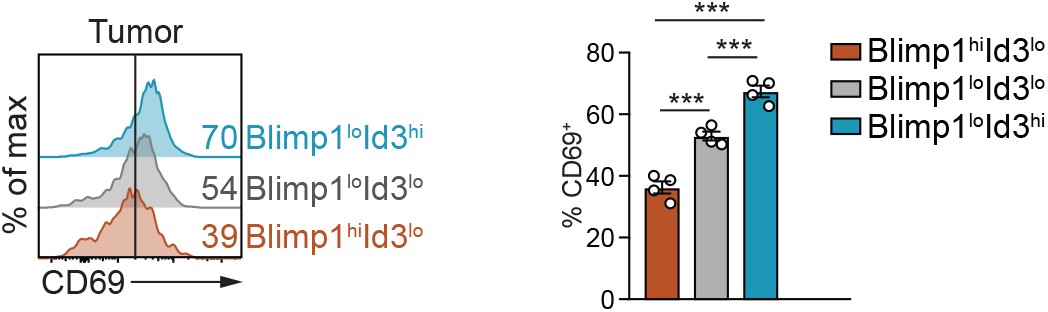
Id3hi Trm-like TIL express elevated levels of CD69. Congenically distinct Id3-GFP/Blimp1-YFP P14 cells were transferred into tumor-bearing mice, and one-week post adoptive transfer, donor cells from tumors were analyzed for CD69 expression. Numbers in plots represent frequency of cells in the indicated gate. All data are from one representative experiment with n=4. Graphs show mean ± SEM; ***p<0.005.

